# A mitochondrial program encodes brain vascular reserve

**DOI:** 10.64898/2026.03.25.714175

**Authors:** Ivan Fan Xia, Dingding Zhang, A. Gulhan Ercan-Sencicek, Tanyeri Barak, Anupama Hemalatha, David G. Gonzalez, Jared Hintzen, Paola Carneiro, Gabriel Baldissera, Marina Carlson, Siyuan Cheng, Fei Han, Xinzhuang Yang, Lu Feng, Nicole J. Lake, Valentina Greco, Murat Günel, Stephanie Debette, Yi-Cheng Zhu, Stefania Nicoli

## Abstract

Brain resilience depends on collateral vessels whose geometries preserve blood flow when primary arteries are perturbed. How these protective vascular architectures are developmentally established remains unknown. Using longitudinal *in vivo* imaging in zebrafish, we show that mitochondrial state in embryonic angiogenic tip cells encodes the topology of basal and surface brain collateral networks. Mechanistically, microRNA-125a establishes the bioenergetic and redox balance of endothelial tip-cell mitochondria through conserved repression of the metabolic regulator *PGC1a*. Disruption of the microRNA-125a–*PGC1a* axis uncouples mitochondrial capacity from redox buffering in developing brain tip cells, redirecting their migration toward sparse, incompletely connected collateral network topologies and increasing adult cerebrovascular vulnerability. Consistent with this mechanism, humans with subclinical cerebrovascular injury exhibit reduced circulating microRNA-125a levels associated with incomplete basal collateral configurations. Together, these findings identify mitochondrial state as an instructive and conserved developmental program that encodes brain vascular reserve.

## Introduction

Collateral circulation provides a vascular “reserve” that can reroute blood flow when primary vessels become occluded ^1–7^. The brain is critically dependent on a continuous blood supply and has two anatomically and functionally distinct collateral networks: the primary collateral circulation named the Circle of Willis (CoW) at the brain base and the secondary collateral circulation, the leptomeningeal vessels distributed across the brain cortical surface. The CoW consists of paired anterior and posterior segments that form an arterial ring ^8,9^, whereas leptomeningeal collaterals form a mesoscopic arterial mesh supplied by CoW arteries and distal branches of its major segments. These vascular architectures are strongly conserved across species, underscoring their evolutionary preservation as protective patterning programs ^10,11^.

Humans display substantial inter-individual variability in both the primary and secondary brain collaterals ^12–16^. For example, 34–68% of asymptomatic individuals exhibit an “incomplete CoW” configuration, characterized by a hypoplastic or absent posterior or anterior segment ^17–20^. Upon cerebral artery occlusion, tissue perfusion can be preserved through compensatory flow rerouting via the CoW and the secondary leptomeningeal (pial) collateral network ^7,11^. Notably, incomplete CoW configurations are associated with reduced leptomeningeal collateralization, vascular vulnerability, and poorer clinical outcomes ^21–23^, indicating that CoW architecture and pial network organization are functionally coupled and together determine cerebrovascular resilience.

Although collateral patterning remodels during vascular occlusion, the genetic background determines baseline brain collateral architecture. In mice, native leptomeningeal collateral density, diameter, and connectivity are established early in development and persist across the lifespan ^24,25^. CoW anatomy has been linked to multiple environmental and genetic factors ^26^, yet whether CoW configuration and pial collateral architecture are coordinated by shared genetic programs in the native (non-occlusive) state remains unknown. A related mechanistic gap is how such shared programs, if they exist, are executed during development to generate coordinated architecture across two anatomically distinct collateral tiers. Defining these mechanisms is essential to understand and ultimately protect cerebrovascular resilience.

Collateral networks assemble early in development through angiogenic sprouting and vessel–vessel connections. These processes are controlled by specialized endothelial states, especially tip cells at the sprout front, which direct migration and connectivity while stalk cells extend and remodel the sprout ^27^. Emerging evidence suggests that tip cells are not interchangeable units but instead adopt organ- and context-specific states early in development ^28,29^. While this specialization has been linked to organ-specific vascularization, whether tip cells encode higher-order vascular patterning across beds remains unknown. Collateral vessels provide an ideal context to address this question: they arise during a narrow developmental window via angiogenesis-like mechanisms, yet display striking inter-individual variability in network geometry and connectivity. Discovering a “patterning tip cell” program that builds native collateral network topologies with functional consequences would provide a new entry point for strategies to enhance vascular resilience in virtually every organ.

Here, we show that the posterior brain collateral architecture, including both primary and secondary networks, is determined by embryonic migrating tip cells through a conserved metabolic program. We show that complete CoW anatomy co-occurs with dense surface collateral networks and identify miR-125a as a key regulator of these coupled topologies in both zebrafish and humans. miR-125a functions during developmental brain angiogenesis to specify a pool of posterior endothelial tip-cells that pattern the angiogenic growth of basal and surface collateral territories through metabolic control. This program operates through miR-125a repression of *pgc1a*-controlled mitochondrial biogenesis and redox state, linking embryonic tip-cell metabolic preferences to multi-tier brain vascular architecture and adult resilience.

## Results

### miR-125a Levels Predict Posterior Brain Collateral Completeness and Vascular Resilience

The extent to which primary and secondary brain collaterals’ structure co-vary under native conditions is undefined in model organisms. We therefore leveraged adult *Tg(kdrl:GFP*)^zn1^ zebrafish, in which endothelial cells are fluorescently labeled, and combined this with microangiography using high–molecular-weight fluorescent dextran to visualize perfused collateral architectures across scales ^30^. The Circle of Willis (CoW) in zebrafish, as in humans contains two anterior cerebral arteries (left and right ACAs) and two posterior cerebral/communicating arteries (left and right PCAs) connected to the posterior basilar artery (BA) (**Figure 1A**). The secondary collateral network, comparable to mammal leptomeningeal or pial vessels, covers the posterior and anterior brain surface. We term these vessels surface brain collaterals (SBCs) (**Figure 1B**).

**Figure 1.**
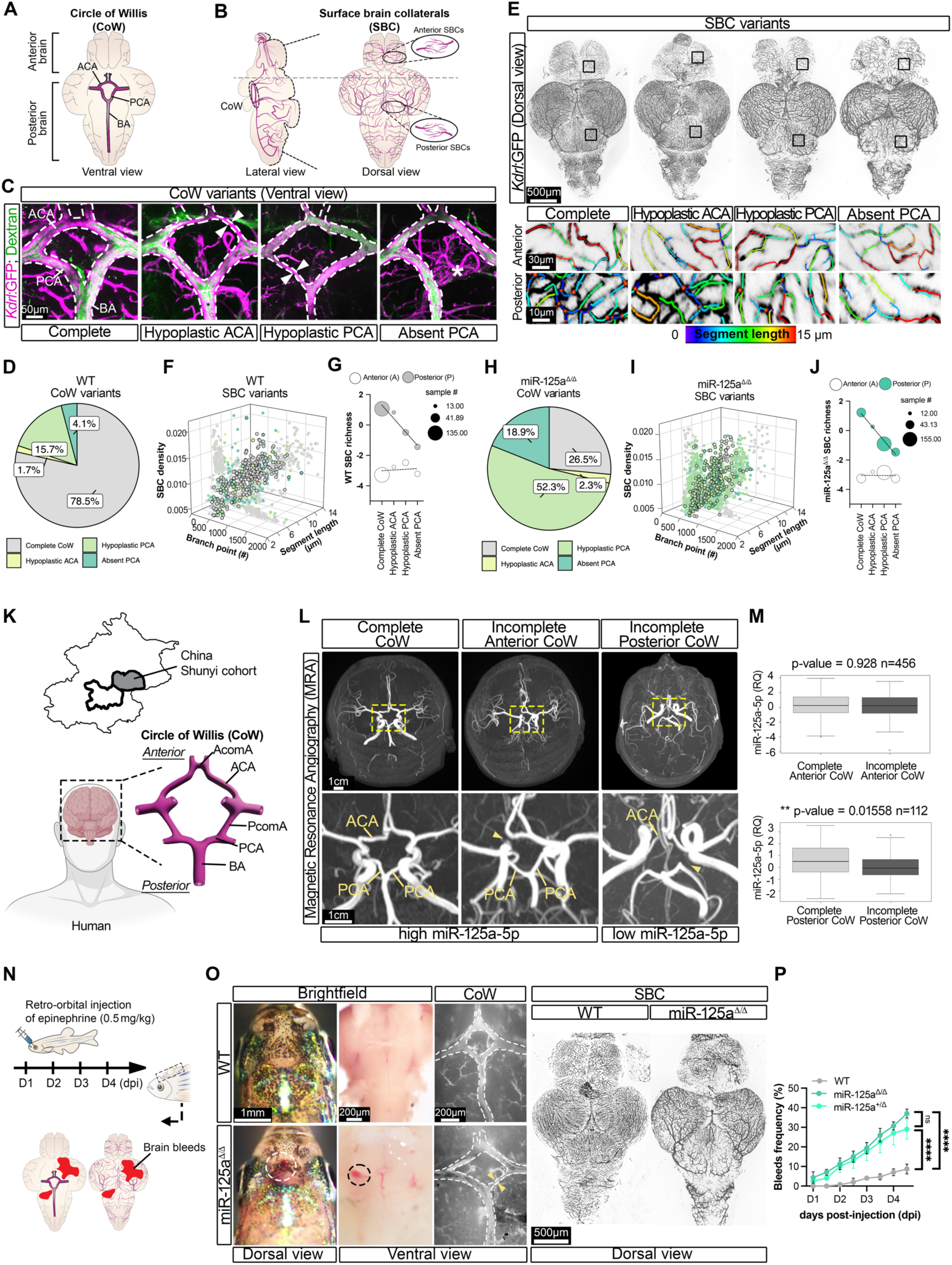
miR-125a levels predict posterior collateral completeness and cerebrovascular vulnerability. (**A**) Schematic of the zebrafish Circle of Willis (CoW) and (**B**) surface brain collaterals (SBCs). Vessel abbreviations below. (**C**) Representative confocal images of adult CoW configurations in wild-type Tg(*kdrl:GFP*)*^zn1^*via intracardiac injection of Tetramethylrhodamine Dextran (2,000 kDa). Magenta = *kdrl:GFP,* green= Dextran, White dash = lumen, Arrows = areas of hypoplasia without flow; asterisk = absent CoW arteries. (**D**) Wild-type adult CoW variant distribution (n = 155 fish). Variants in the AComA not quantified due to biases that brain dissection can infer on these vessels (see STAR Methods). (**E**) Representative confocal images of Tg(*kdrl:GFP*)*^zn1^*in C of anterior and posterior SBC networks among each CoW variant, box=magnified brain zones. Individual vessel segments color-coded by branch segment length. Scale bars as shown. (**F**) 3D scatter plot of SBC morphometric parameters including density normalized to brain area (μm/μm^2^, z-axis), branch point number (#, x-axis), and mean branch segment length (μm, y-axis) in wild-type. Each dot represents the morphometric measurements of the anterior or posterior SBC network from a single wild-type fish; therefore, individual fish may contribute up to two data points (n = 155 fish). (**G**) Correlation between anterior or posterior SBC richness and CoW configurations in wild-type. SBC richness via principal component analysis (see STAR Methods). Pearson correlation analysis shows anterior r= 0.161, p= 0.839; posterior r= −0.972, p= 0.028. (**H**) Adult miR-125a^Δ/Δ^ CoW variant distribution (n = 190 fish). (**I**) 3D scatter plot of SBC morphometric parameters in miR-125a^Δ/Δ^ zebrafish, analogous to panel F (n = 190 fish). (**J**) Correlation between anterior or posterior SBC richness and CoW configuration in miR-125a^Δ/Δ^. Pearson correlation analysis shows anterior r= −0.036, p= 0.964; posterior r= −0.991, p= 0.008, n=190 fish (**K**) Schematic of human CoW and geographic localization of the Shunyi community-based cohort. (**L**) Representative magnetic resonance angiography (MRA) images from individuals with complete CoW, incomplete anterior CoW, or incomplete posterior CoW. Magnified view below, Scale bars as shown. (**M**) RT-qPCR quantification of miR-125a-5P from peripheral blood samples, normalized to housekeeping gene, *U6* (RQ: relative quantification, calculated as 2^(-ΔΔCT)^) for individuals with complete and incomplete anterior (p-value= 0.928, n=456) or posterior (p-value= 0.01558, n=112) CoW. Log transformation and Wilcoxon signed rank test were performed after age and sex matched pairs. (**N**) Acute hemodynamic stress model in adult zebrafish (6-12 months old) via twice-daily retro-orbital epinephrine injections for four days. (**O**) Representative dorsal brightfield images of adult fish heads (left), alongside brightfield brain base (middle) and the corresponding fluorescent images of dissected CoW configuration (right, *kdrl:GFP*) in ventral view, and SBC network in dorsal view. Dashed circle=bleed event. Dashed line = CoW vessels. Arrow=hypoplastic arteries. (**P**) Quantification of brain bleed frequency in adult WT, miR-125a^+/i1^ and miR-125a^ι1/i1^. Each group consisted of 30 fish from three independent experiments, dots represent average over all experiments. Ordinary two-way ANOVA test at D4 (p=6.71e-9(WTvs125a^Δ/Δ^), 8.61e-8(WTvs125a^+/Δ^), 0.75(125a^+/Δ^vs125a^Δ/Δ^). All quantifications are presented as mean ± s.e.m. unless otherwise noted. Statistical significance is indicated: ns, not significant, p:: 0.05; *p< 0.05; **p< 0.01; ***p< 0.001; ****p< 0.0001. All data points are reported in Table S1 Sheets 1, 2, 3, 17. Abbreviations: CoW= the Circle of Willis; SBC= surface brain collaterals; ACA= anterior cerebral artery; PCA= posterior cerebral/communicating artery; BA= basilar artery.

We found that 78.5% ± 5.7% of individuals in our standard adult wild-type population possessed a complete CoW anatomy (**Figures 1C, 1D, S1A and S1B**). We also detected incomplete CoW configurations, including hypoplastic ACAs (1.7% ± 3.0%) and hypoplastic or absent PCAs (19.8% ± 3.9%) (**Figures 1C and 1D**). Hypoplastic vessels were defined as having a diameter less than 50% of the contralateral vessel. Posterior CoW anatomical variants (hypoplastic or absent PCA) were strongly associated (p<0.0001) with overall CoW incompleteness, whereas body weight, sex, and anterior CoW variants were not (**Figure S1C**).

Next, we analyzed zebrafish secondary anterior and posterior SBCs by quantifying multiple vascular parameters, including vessel density, branch-point count, and segment length. These features were integrated into a single composite metric, termed the richness score (see STAR Methods), which captures the overall structural complexity of the vessel network. We found that SBCs exhibited substantial heterogeneity across individuals and between anterior and posterior brain regions: some individuals formed sparse, minimally interconnected networks, while others developed dense, highly branched patterns with anterior networks being generally more sparse than posterior ones (**Figures 1E, 1F and S1D)**. Strikingly, heterogeneity in posterior—but not anterior—SBCs was not random but correlated with CoW anatomy, with hypoplastic or absent PCA variants emerging as the strongest predictors of reduced SBC richness (**Figures 1C, 1E, 1G and S1E**). Thus, native posterior SBC network architecture is tightly interdependent with the posterior configuration of the CoW.

We next hypothesized that shared mechanisms co-establish posterior native architectures and set collateral reserve. Because microRNAs regulate the variability of specific vascular traits ^31^, we surveyed CoW anatomy across a panel of microRNA mutants previously generated for vascular phenotypic screening ^32^ to identify candidate regulators (**Figure S1F**). We found that loss of microRNA-125a (miR-125a^Δ/Δ^) markedly increased the frequency of incomplete CoW, with 71.2% ± 7.6% of adult animals exhibiting incomplete posterior configurations (**Figures 1H and S1F**). These defects most commonly presented as hypoplastic (52.3% ± 5.2%) or absent (18.9% ± 4.1%) PCAs with ACA configuration comparable to wild-type (2.3% ± 1.3%) (**Figures 1H, S1G and S1H**). miR-125a^Δ/Δ^ animals displayed anterior and posterior SBC variability comparable to wild-type, but individuals with incomplete PCA configurations exhibited reduced vessel density, fewer branch points, and remodeling of segment architecture only in the posterior brain region (**Figures 1I, 1J**, **S1I and S1J**). Hence, the increased incidence of incomplete CoW remained strongly associated with reduced posterior, but not anterior, SBC network richness (**Figure 1J**). miR-125a^Δ/Δ^ animals exhibited no overt morphological abnormalities and were viable and fertile (**Figure S1K**). Together, these findings establish miR-125a as a determinant of collateral completeness across tiers, linking its loss to incomplete posterior primary and secondary collateral architectures.

We next asked whether this relationship extends to humans ^7,22,33^. MicroRNAs are highly stable in the circulation and are considered long-term biomarkers of vascular dysfunction ^34–37^. We therefore analyzed circulating blood miR-125a levels and CoW anatomy in a cohort from the Shunyi study, in which individuals with incomplete CoW collateral variants frequently display radiological markers of covert vascular brain injury ^38–40^ (**Figures 1K and S2A-S2D**). Strikingly, we found that individuals with complete posterior CoW collaterals have significantly higher circulating mature miR-125a levels than those with incomplete posterior variants, whereas no significant differences were detected across anterior variants (**Figures 1L and 1M**). Altogether, these data support a conserved association between miR-125a levels and incomplete posterior cerebrovascular collateral architecture.

Given that in the Shunyi cohort incomplete CoW is associated with covert vascular injury, including cerebral microbleeds, we tested whether the incomplete CoW and SBC density observed in miR-125a^Δ/Δ^ zebrafish associates with increased susceptibility to stress-induced vascular bleeds. We exposed miR-125a^Δ/Δ^ adults to a hypertensive-stress challenge with repeated retro-orbital injections of epinephrine over four days ^30^ (**Figure 1N**). Indeed, miR-125a^Δ/Δ^ and miR-125a^+/Δ^ displayed substantially increased brain bleeding, with a ∼4-fold higher bleed frequency compared to wild-type by day 4 (**Figures 1O and 1P**). Moreover, miR-125a^+/Δ^ displayed both CoW and SBC anatomical deficits (**Figure S2E),** indicating that partial downregulation of miR-125a is sufficient to compromise posterior cerebrovascular collateral architecture and resilience to vascular stress (**Figures 1P and S2E-S2G**).

Together, these findings identify that the architectural patterning of posterior brain collaterals is regulated by miR-125a level with functional consequences for cerebrovascular robustness from zebrafish to humans.

### Endothelial miR-125a Directs the Development of Complete Brain Collateral Variations

Leptomeningeal vascular architecture is established during embryonic development ^24,41^. To determine if the coordinated patterning of primary posterior collaterals (PCA within the CoW) and associated secondary posterior networks (SBCs) is also established during embryogenesis, we examined zebrafish at 5 days post-fertilization (dpf), when both the primordial structure of the posterior CoW and SBCs (Central Arteries -CtAs-) can be recognized ^42^ (**Figure 2A).** Notably, miR-125a^Δ/Δ^ larvae displayed a markedly higher frequency of hypoplastic or absent posterior PCA segments (40.9% ± 8.1%) compared with wild type (**Figures 2B, 2D, 2E and S3A**). Furthermore, posterior CtA richness was diminished in mutant larvae and inversely correlated with posterior PCA segment variation in both genotypes, consistent with coordinated disruption of posterior collateral architecture (**Figures 2C, 2F and S3B**).

**Figure 2.**
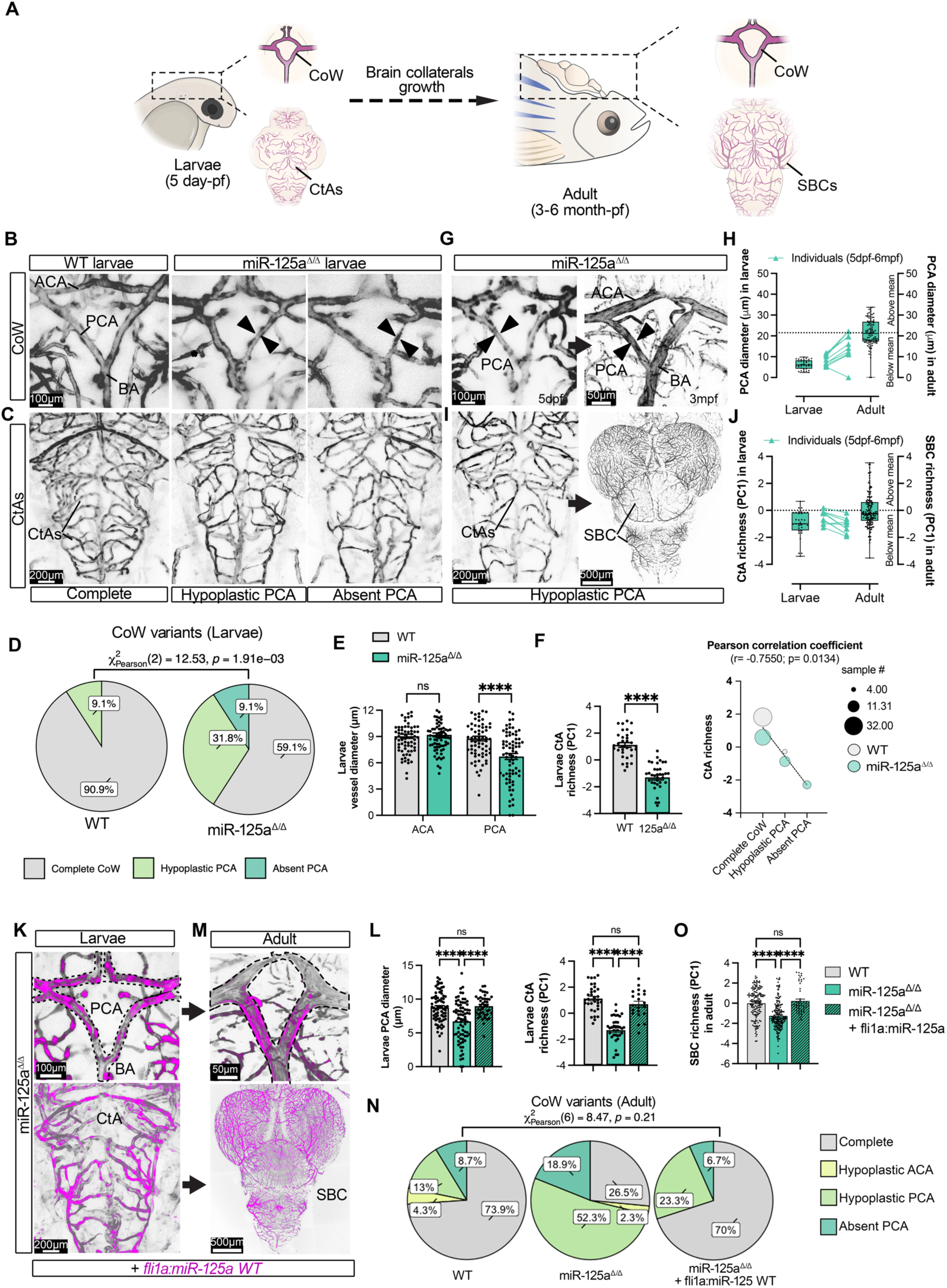
Developmental origin of primary and secondary brain collateral architectures. **(A)** Schematic overview of brain collateral development across larval and adult stages. Abbreviations as in Fig. 1 and below, superficial central arteries (CtAs) are considered larval SBCs. **(B)** Representative dorsal confocal images of larval CoW anatomy in wild-type (WT) and miR-125a^Δ/Δ^ zebrafish at 5 days post fertilization (dpf). Arrowheads= hypoplastic (left, defined as less than 50% width of the contralateral vessel) or absent (right) where PCA segment was found disconnected from ACA arteries. Scale bars shown. **(C)** Representative dorsal confocal images of larval CtA anatomy in WT and miR-125a^Δ/Δ^ zebrafish at 5 dpf. Scale bars shown. (**D**) Distribution of larval CoW variants in WT and miR-125a^Δ/Δ^ animals at 5 dpf, classified as complete, hypoplastic PCA, or absent PCA. (Pearson’s χ² test of independence (χ²(2) = 12.53, ** p=1.91e-3, Cramer’s V =0.35, n = 36 WT larvae and 38 miR-125a^Δ/Δ^). **(E)** Quantification of larval ACA and PCA CoW vessel diameter. Each dot represents a left or right vessel per embryo, n=72 WT vessels and 76 miR-125a^Δ/Δ^ vessels. Multiple unpaired t test, ACA, p=0.625, PCA, p=9.754e-6. (**F)** Quantification of larval CtA richness score in WT and miR-125a^Δ/Δ^ zebrafish. Each dot represents one larva, n = 36 WT larvae and 38 miR-125a^Δ/Δ^ larvae. Mann–Whitney test, p=1.55e-11; Correlation analysis for CtA richness with CoW variants in WT and miR-125a^Δ/Δ^ zebrafish. (Right) Pearson correlation analysis for both groups, r = −0.7550, p = 0.0134. **(G)** Representative longitudinal confocal images of larvae (left) and adult (right) miR-125a^Δ/Δ^ zebrafish CoW anatomy, scale bars shown. **(H)** Paired analysis of PCA diameter measured in the same individual at larval (5 dpf) and adult (6 mpf) stages. Each line represents one larvae traced to adulthood (n= 5), and boxplots show vessel diameters (dots) from n= 22 miR-125a^Δ/Δ^ larvae and 50 adults. Dashed line= adult mean value. **(I)** Larval CtA (left) and adult SBC (right) anatomies. **(J)** Paired analysis of CtA richness score measured in individual animals at larval and adult stages, as in H. **(K)** Representative images of *fli1a:miR-125a-mCherry* expression in miR-125a^Δ/Δ^ larval CoW (top) and CtA (bottom) vessels. Scale bars as shown. **(L)** Quantification of larval PCA diameter and richness. (left) Each dot represents one PCA vessel, and (right) each dot represents one larva, n = 36 WT larvae, 38 miR-125a^Δ/Δ^ and 22 *fli1a* rescue, one-way ANOVA, p=0.0002, p=2.00e-15. (**M**) Adult CoW and SBC configurations from larvae in K. Scale bars shown. (**N**) Distribution of adult phenotype in *fli1a* rescue animals. Pearson’s χ² test of independence revealed no significant difference among WT, miR-125a^Δ/Δ^, and with *fli1a* miR-125a rescue groups (χ²(6) = 8.47, p = 0.21; ns, Cramer’s V=0.28, n = 155 fish WT, 190 miR-125a^Δ/Δ^ and 55 *fli1a* rescue adult), and WT and *fli1a* rescue adults were indistinguishable in CoW configuration (χ²(3) = 2.13, p = 0.55, ns, Cramer’s V=0.00). Additional comparisons in S2.1I. **(O)** Quantification of adult SBC richness among WT, miR-125a^Δ/Δ^, and *fli1a* rescue groups. Each dot represents one fish from n = 155 WT, 190 miR-125a^Δ/Δ^ and 55 *fli1a* rescue adults. One-way ANOVA, p=9.22e-17. All quantifications are presented as mean ± s.e.m. unless otherwise noted. Statistical significance is indicated: ns, not significant, p:: 0.05; *p< 0.05; **p< 0.01; ***p< 0.001; ****p< 0.0001. All data points are reported in Table S1 Sheets 4, 5. Abbreviations: CtA, central artery; dpf, days post-fertilization; mpf, months post fertilization.

To test whether this early defect in vascular architecture leads to the collateral outcomes in adults, we performed longitudinal live imaging in individual animals, tracking disruptions in PCA–CtA architecture from larval stages into adulthood (**Figure 2A**). Importantly, miR-125a^Δ/Δ^ mutant embryos have no major defect: trunk vasculature, red blood cell circulation, and heart rate were indistinguishable from wild-type controls (**Figures S3C-S3E**). Remarkably, the coordinated defects in PCA caliber and CtA network complexity at 5 dpf persisted into adulthood, manifesting as reductions in posterior CoW completeness and secondary collateral richness (**Figures 2G-2J).** These data strongly suggest that miR-125a^Δ/Δ^ adult configurations are already present at 5 dpf.

Collateral vessels form within a complex multicellular environment, so defects could reflect non-endothelial influences; however, miR-125a was originally identified as an early endothelial-enriched microRNA ^31^ (**Figures S1F, S3F and S3G**). To test whether the coordinated PCA–CtA defects arise from an endothelial-autonomous requirement for miR-125a, we stably re-expressed wild-type miR-125a gene specifically in endothelial cells via a pan *fli1a* endothelia promoter, creating an miR-125a^Δ/Δ^ *fli1a:miR-125a–mCherry* transgenic endothelial rescue model (**Figures 2K and 2M**). Importantly, endothelial-specific miR-125a rescued posterior cerebrovascular architecture, simultaneously normalizing PCA diameter and posterior CtA richness at larval stages (**Figures 2K, 2L and S3H**). Tracking of rescued animals demonstrated sustained correction of adult CoW configuration and secondary collateral network complexity (**Figures 2M-2O, S3I and S3J**).

Together, these findings demonstrate that endothelial miR-125a coordinates the coupled development of primary and secondary posterior brain collateral architectures.

### miR-125a Is Required to Coordinate Territorial Brain Sprouting during Embryonic Angiogenesis

Since the incomplete collateral architecture in miR-125a^Δ/Δ^ is already evident at larval stages, we reasoned that it can be traced back to aberrant endothelial cell behaviors during the establishment of the nascent structure. In wild-type embryos, posterior CoW arteries and CtAs arise through coordinated angiogenic sprouting from a shared pool of endothelial cells emerging from the primordial hindbrain channel (PHBC) ^43^. Beginning at ∼26–28 hpf, endothelial sprouts from the PHBC give rise to the posterior communicating arteries (EC^PCA^) and basilar artery (EC^BA^), which subsequently fuse at the midline by ∼32 hpf. In parallel, at ∼28 hpf CtA endothelial cell (EC^CtA^) segments extend dorsally and connect to the basilar artery approximately 16 hours later (**Figure 3A**).

**Figure 3.**
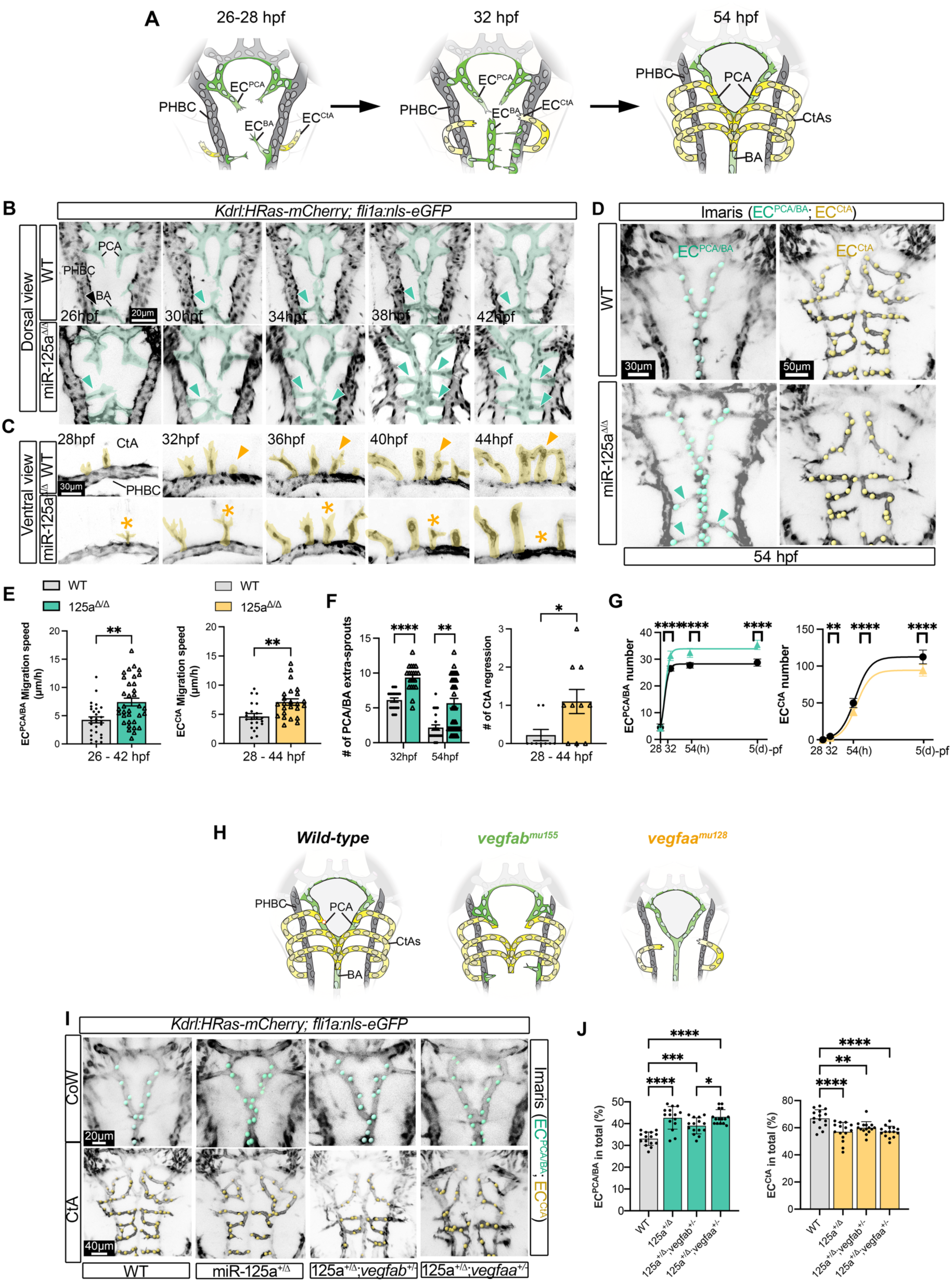
miR-125a deficiency induces divergent angiogenic behaviors in posterior brain collaterals. **(A)** Schematic overview of posterior brain angiogenesis in zebrafish embryos. PCA and BA arteries (CoW, green), as well as CtA (yellow) form from PHBC sprouts which subsequently supply distinct vascular territories. Abbreviations below. **(B, C)** Representative time-lapse images of dorsal CoW angiogenesis (B) and ventral CtA angiogenesis (C) in *Tg(kdrl:HRAS-mCherry; fli1a:nls-eGFP)* embryos (26 to 44 hpf) of both wild-type (WT, top, Videos S1 and S2) and miR-125a^Δ/Δ^ (bottom, Videos S3 and S4). Arrows= sprouting, Asterisk= sprout regression. Scale bars as shown. **(D)** Representative Imaris-labeled endothelial nuclei from *Tg(kdrl:HRAS-mCherry; fli1a:nls-eGFP)* in WT and miR-125a^Δ/Δ^ at 54 hpf. Posterior CoW (PCA/BA in green) and CtA in yellow. Arrowheads indicate abnormal extra PCA/BA-associated sprouts in mutants. Scale bars as shown. **(E)** Quantification of endothelial migration speed during PCA/BA (left) and CtA sprouting (right) in WT and miR-125a^Δ/Δ^ embryos. Each data point represents one migrating endothelial cell, (n=4-5 cells/timelapse). For PCA/BA sprouts, n= 5 timelapses each genotype and for CtA sprouts, n= 4 WT timelapses and 6 miR-125a^Δ/Δ^ timelapses. Mann–Whitney test, p=0.0023, 0.0027. (**F**) Quantification of extra sprouts in CoW (left) and CtA regression events (right) in WT and miR-125a^Δ/Δ^. For extra sprouts in CoW, each dot represents one embryo from n = 20 embryos each genotype at 32hpf; for CtA regression, each dot represents one CtA vessel from n= 9 timelapses each genotype. Mann–Whitney test, p=3.03e-7, 0.038. **(G)** Average of endothelial cell (EC) number in PCA/BA vessels (left) and CtA (right) over time. For 32hpf n= 25 embryos each genotype, 54 hpf n= 20 each genotype, and 5 dpf n=13 each genotype. Multiple Mann–Whitney test for 32, 54, 5dpf respectively, PCA/BA, p=1.14e-5, 1.96e-5, 1.07e-4 and CtA p=0.0080, 2.57e-8, 2.70e-7. **(H)** Schematics illustrating *vegfa* isoform-specific contributions to posterior brain angiogenesis: *vegfab* primarily contributes to PCA/BA and *vegfaa* predominantly supports CtA angiogenesis. **(I)** Representative Imaris-labeled endothelial nucleus from *Tg(kdrl:HRAS-mCherry; fli1a:nls-eGFP)* for posterior CoW (green) and CtA (yellow) vessels in WT, miR-125a^+/Δ^, miR-125a^+/Δ^;*vegfab^+/Δ^*, miR-125a^+/Δ^;*vegfaa^+/Δ^* embryos. (**J**) Bar plot represents the percentage of ECs residing in PCA/BA or CtA relative to the total number of posterior brain ECs. Each dot represents one embryo from n=15 embryos each genotype. One-way ANOVA, p=4.021e-9, 1.842e-5. All quantifications are presented as mean ± s.e.m. unless otherwise noted. Statistical significance is indicated: ns, not significant (not shown), p:: 0.05; *p< 0.05; **p< 0.01; ***p< 0.001; ****p< 0.0001. All data points are reported in Table S1 Sheets 6, 7. Abbreviations: PHBC=primordial hindbrain channel; CoW=Circle of Willis; BA=basilar artery; PCA=posterior cerebral/communicating artery; CtA=central artery; hpf=hours post-fertilization; EC= endothelial cell.

We performed live imaging of *Tg(fli1:negfp)^y7^;* (*kdrl:hras-mcherry*)^s896^ embryos to track single endothelial cell behaviors. In wild type, both EC^CtA^ and EC^PCA/BA^ sprouting and migration were well-coordinated and resolved by 54 hpf as expected ^42,43^ (**Figures 3B-3D; Videos S1 and S2**). In contrast, in miR-125a^Δ/Δ^ embryos, although EC^PCA/BA^ and EC^CtA^ initiated angiogenic sprouting from PHBC on schedule, they exhibited profoundly dysregulated behaviors across both posterior CoW and cerebral artery territories (**Figures 3B-3D; Videos S3 and S4**). PHBC-derived endothelial cells generated sprouts and migrated at abnormally high speeds, producing aberrant trajectories that spanned both posterior (BA/PCA) and cerebral artery domains (**Figures 3B, 3C and 3E**).

Unexpectedly, despite this shared hyper-migratory behavior, subsequent remodeling diverged in a territory-specific manner. BA and PCA sprouts failed to properly remodel, retaining excess endothelial cells that persisted through 5 dpf, forming narrowed, hypoplastic vessel segments (**Figures 3F, 3G, S4A and S4B**). In contrast, CtA vessels were unstable: although initiated normally, EC^CtA^ frequently underwent regression within 5 hours of sprout emergence, with one to two CtA vessels retracting back toward the PHBC (**Figures 3C and 3F**). As a result, CtA networks contained fewer endothelial cells per vessel by 54 hpf and remained reduced through 5 dpf (**Figures 3D, 3G and S4A**). Together, these findings show that loss of miR-125a uncouples deployment and stabilization of PHBC-derived endothelial cells, driving excessive but poorly remodeled posterior CoW growth alongside instability and loss of central artery segments and density.

The establishment of posterior CoW and CtA networks has been attributed to the endothelial response to regional *vegfa* isoform gradients ^44^, with *vegfab* and *vegfaa* preferentially guiding posterior CoW arteries and CtA sprouting formation, respectively (**Figure 3H**). Therefore, we considered that the divergent sprouting of miR-125a^Δ/Δ^ EC^PCA/BA^ vs EC^CtA^ phenotypes might reflect altered vegfa-signaling. However, we did not detect changes in *vegfab* and *vegfaa* expression in miR-125a^Δ/Δ^ relative to wild-type (**Figures S4C-S4F**). Heterozygous loss-of-function mutant *vegfab^mu1^*^55^ or *vegfaa^mu1^*^28^ embryos in a wild-type background displayed the expected CtA and BA phenotypes ^44^ (**Figures S4G and S4H)** while introducing these alleles into the miR-125a^+/Δ^ background did not alter posterior arteries’ defects beyond what was already observed in miR-125a^+/Δ^ alone (**Figures 3I and 3J**). These findings suggest that the miR-125a mutant phenotype is not driven by altered *vegfa*-signaling during posterior artery angiogenesis.

Together these data demonstrate that miR-125a is dictating distinct endothelial-intrinsic mechanisms within posterior CoW and CtA angiogenesis.

### miR-125a Instructs Tip-Cell Specialization to Build Brain Vascular Territories

The growth of a vascular network is determined by tissue-specific endothelial tip cells with distinct angiogenic behaviors ^28^. We hypothesized that miR-125a level might differentially regulate CoW and CtA sprouting behaviors via tip cell specification. To test this hypothesis, we first turned to human vascular organoids (hVOs), which model human vascular development, including developmental angiogenesis and tip cell behavior ^45,46^ (**Figure 4A**). We established hVOs from induced pluripotent cells, and perturbed miR-125a expression during the mesodermal stage, prior to vascular specification, by lentiviral delivery of an miR-125a inhibitor co-expressing mCherry (miR-125a^inh^) or a scrambled mCherry control (scramble) (**Figures 4A-C**). miR-125a expression showed ∼50–60% reduction at both vascular unit (D5) and hVO stages (D21) (**Figure 4D**). In both conditions, CD31+ endothelial networks with lumen-forming tip cells developed, consistent with robust angiogenesis in hVOs ^45^ (**Figure 4C**). However, CD31+ and mCherry+ cells carrying the miR-125a^inh^ were significantly enriched in the angiogenic front with characteristic tip-cell morphology (extended filopodia from sprouting tips) and lower ColIV staining in comparison to control cells (**Figures 4C** and **4E)**.

**Figure 4.**
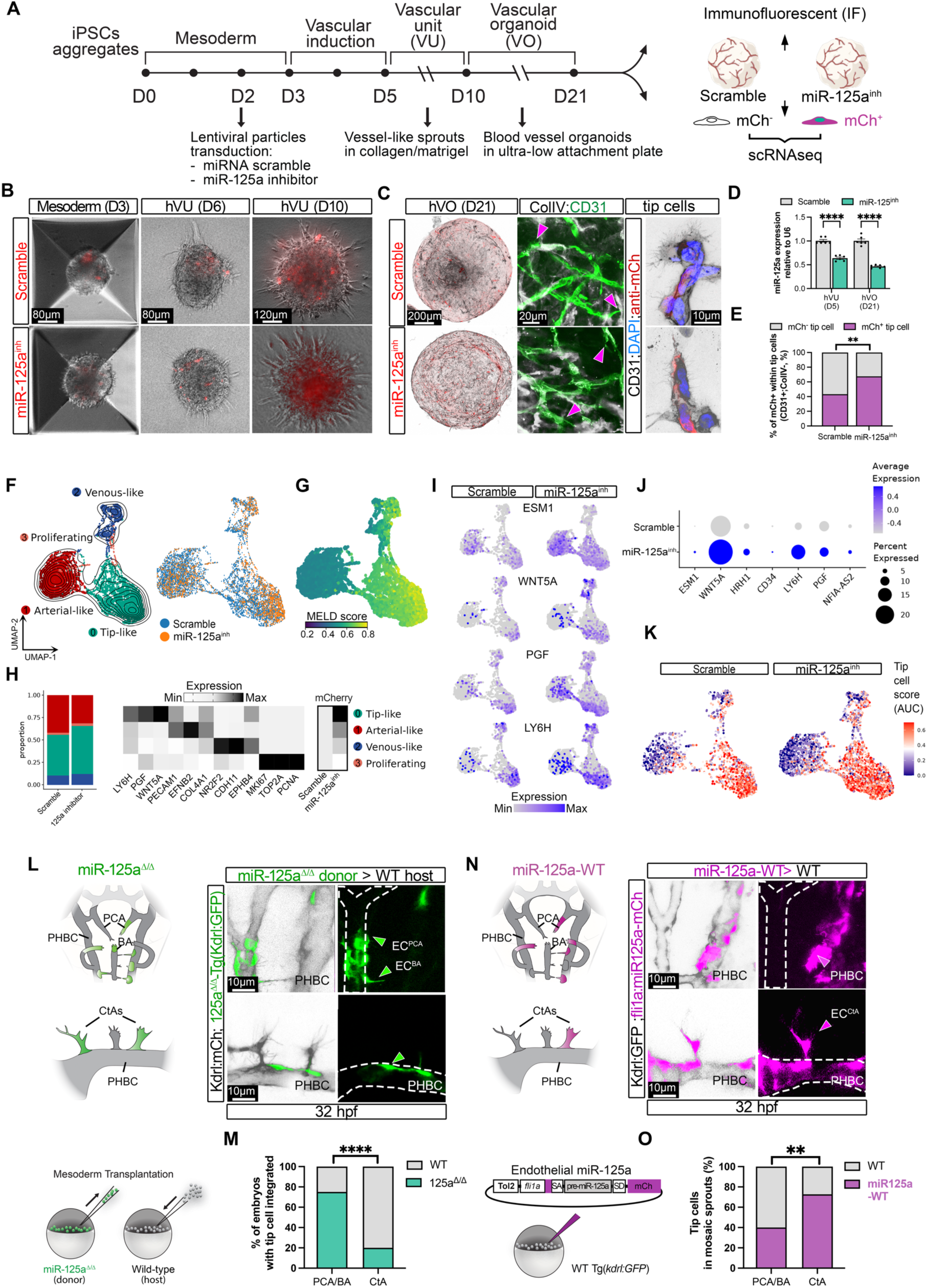
miR-125a regulates the specification of basal and surface brain tip cells. **(A)** Schematic of human induced pluripotent stem cells (iPSCs) differentiation to vascular unit (hVU) and organoid (hVO) and experimental design. Lentiviral particles encoding a miR-125a inhibitor (miR-125a^inh^) or scrambled control, both co-expressing mCherry, were introduced during the mesodermal stage. **(B)** Representative brightfield images of iPSC mesoderm (D3) and hVU (D6, D10) transduced with scramble or miR-125a^inh^. Scale bars shown. **(C)** (Left) Representative image of hVO (D21) with scramble or miR-125a^inh^. (Middle) Immunofluorescent stain for CD31+ endothelial cells (green) and collagen IV (gray). Arrows=CD31+ and ColIV-cells with tip-cell morphology at the angiogenic front. (right) Representative mCherry+ transduced hVO tip cells. Scale bars as shown. **(D)** Quantification of miR-125a expression in infected hVU (D5) and hVO (D21), measured by RT–qPCR. Each dot represents three technical replicates derived from 2 independent experiments. Multiple unpaired t test, p=1.67e-6 (D5), p=1.04e-5 (D21). **(E)** Quantification of the relative contribution of mCherry+ cells to CD31+ ColIV-tip-cell regions in scramble and miR-125a^inh^ organoids. n= 15 hVOs per group, derived from three independent differentiation experiments. Two-tailed Mann–Whitney test, p=0.0021. **(F)** UMAP projection of hVO scRNA-seq, showing endothelial arterial, venous, proliferating, and tip cell subclusters (left) and treatment condition (right). Total n=8,125 cells, see also Figure S5. **(G)** MELD score overlay reflecting miR-125a–dependent transcriptional perturbation. **(H)** (left) Proportional distribution of scramble and miR-125a^inh^ cells across endothelial subclusters. (right) Heatmap of representative endothelial subtype marker genes in scramble and miR-125a^inh^ conditions. **(I)** Tip cell marker gene expression in scramble and miR-125a^inh^ conditions. **(J)** Dot plot of endothelial and tip-cell marker gene average expression. Dot size=percentage of cells. **(K)** UMAP visualization of tip cell scores for scramble and miR-125a^inh^ cells. Gene sets were defined based on previous iPSC-hVO scRNAseq publication (*45*), and scores were computed using AUCell (AUC-based single-cell gene set scoring). **(L)** Schematic and representative images of transplantation experiments in zebrafish embryos. miR-125a^Δ/Δ^ kdrl:GFP+ donor cells (green) were transplanted into wild-type kdrl:hRAS-mCherry hosts (gray) at the mesoderm stage. Dashed line=vessel boundary. Abbreviations below. **(M)** Quantification of the percentage of embryos exhibiting miR-125a^Δ/Δ^ donor cells at tip-cell positions within posterior CoW (PCA/BA) and CtA sprouts. Fisher’s exact test, total n= 54 embryos, p=8.33e-5. **(N)** Schematic and representative images of endothelial-specific miR-125a gain-of-function mosaic experiments using (*fli1a:miR125a-mCherry*) injection in WT embryos. Donor cell (magenta) localization within posterior CoW and CtA sprouts (gray). Dashed line= vessel boundary. **(O)** Quantification of the percentage of tip-cell positions occupied by miR-125a over-expressing cells in posterior CoW (PCA/BA) and CtA sprouts Fisher’s exact test, total n= 44 embryos, p=0.0039. All quantifications are presented as mean ± s.e.m. unless otherwise noted. Statistical significance is indicated: ns, not significant, p:: 0.05; *p< 0.05; **p< 0.01; ***p< 0.001; ****p< 0.0001. All data points are reported in Table S1 Sheets 8, 9, 12. Abbreviations as in Figure 3.

We next used single-cell RNA sequencing (scRNA-seq) to characterize cellular composition across miR-125a^inh^ and scramble-transduced hVOs. Cell clustering analysis resulted in nine molecularly distinct groups, which were annotated based on cluster marker genes as done previously ^45^ (**Figure S5A-S5D, Table S1 Sheet 8**). Endothelial cell groups were further subclustered to identify arterial, venous, and tip-cell populations using human defining markers (**Figure 4F**). miR-125a–deficient cells were present across all endothelial subclusters; however, relative to controls, they were enriched within the tip-cell cluster and displayed the strongest transcriptional differences (**Figures 4F-4H**). Furthermore, loss of miR-125 increased the expression of tip-cell markers, reflected by an increased tip cell score (**Figures 4I-4K**). Altogether, these findings suggest that miR-125a restrains tip-cell molecular specification.

To test whether miR-125a–dependent tip-cell specification drives posterior CoW and CtA growth *in vivo*, we performed mosaic transplantation assays in zebrafish embryos. We introduced miR-125a^Δ/Δ^*-kdrl:GFP* mesodermal cells into wild-type *kdrl:mCherry* hosts, and observed their preferential localization to tip positions in posterior CoW vessels, with reduced contribution to CtA sprouts (**Figures 4L and 4M**). In contrast, *fli1a:miR125a-mCherry* mesodermal cells overexpressing miR-125a were enriched at CtA tips and depleted from posterior CoW sprouts in wild-type hosts (**Figures 4N and O**). Importantly, similar perturbations of miR-125a did not alter tip cell positioning in trunk intersegmental vessels (**Figures S5E and S5F**), indicating that this bias is specific to brain collateral territories.

Together, these data demonstrate that miR-125a–driven tip-cell specialization instructs the angiogenesis of both posterior basal and surface brain collateral networks.

### *pgc1a*-Mediates miR-125a–Dependent Brain Tip-Cell Specialization

To explore how miR-125a drives tip cell-specialization across brain territories, we first isolated brain endothelial cells from both miR-125a^Δ/Δ^ and wild-type embryos during the formation of the CoW and CtA (∼26-28 hpf) (**Figure 5A**). Given that miRNAs serve to repress target transcripts, we conducted bulk RNA-seq to elucidate genes that were de-repressed in mutant versus wild-type endothelial cells (**Figure 5B** and **Table S1, Sheet 10**). Notably, the peroxisome proliferator-activated receptor gamma coactivator-1 alpha, or *pgc1a* ^47^ (also referred to as *ppargc1a* in zebrafish), was the most significantly upregulated predicted miR-125a target-gene at ∼26 hpf (**Figure 5B**). *pgc1a* contains two conserved miR-125a binding sites in zebrafish and human 3′UTRs, and we found that the wild-type miR-125a mature sequence robustly repressed *GFP–pgc1a-3′UTR* compared with the *mCherry–3′UTR* control lacking miR-125a binding sites in zebrafish embryos (**Figures S6A-S6C**). These data support a model in which miR-125a directly binds the *pgc1a 3′UTR* to mediate post-transcriptional repression.

**Figure 5.**
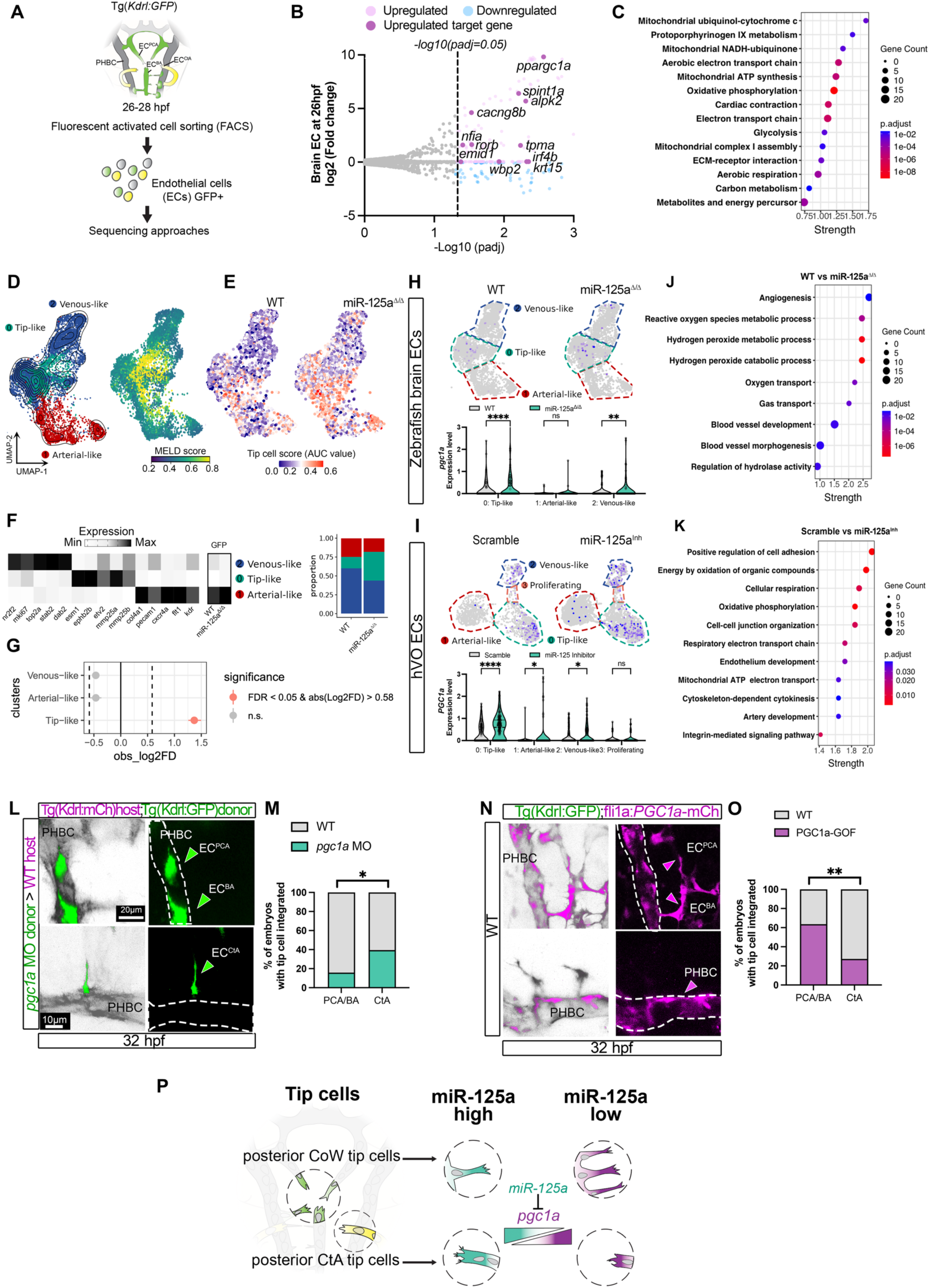
*pgc1a* is a miR-125a target gene and controls tip cell specification in posterior brain angiogenesis. **(A)** Schematic of fluorescence-activated cell sorting (FACS) for bulk and single cell-RNAseq analyses of zebrafish brain endothelial cells at early sprouting angiogenesis (26-28hpf) using *Tg(kdrl:GFP)* embryos. **(B)** Volcano plot of differentially expressed genes (DEGs) in endothelial cells isolated from miR-125a^Δ/Δ^ versus wild-type (WT) embryos at 26hpf and analyzed for bulk RNA-seq. Significant DEGs (padj ≤ 0.05) are shown (up, magenta; down, blue) and upregulated miR-125a target genes (magenta solid) are highlighted. n=five biological replica per genotype, total 9,175 cells. **(C)** Top 14 Gene Ontology and KEGG pathway terms (padj ≤ 0.05) from upregulated genes in 26-28hpf miR-125a^Δ/Δ^ endothelial cells **(D)** (left) UMAP projection of 26-28hpf zebrafish brain endothelial scRNAseq, subclustered as arterial, venous, and tip-cell based on canonical marker expression ^45^. (Right) MELD score overlay reflecting miR-125a–dependent transcriptional perturbation. Total n=9,175 cells, see also Figure S6. **(E)** UMAP visualization of tip-cell gene signature score using area-under-the-curve (AUC) analysis in WT and miR-125a^Δ/Δ^. **(F)** (left) Heatmap showing expression of representative marker genes across endothelial subtypes: arterial, venous, and tip cell-like clusters in WT and miR-125a^Δ/Δ^. (right) Proportional distribution of miR-125a^Δ/Δ^ cells across subclusters. **(G)** Quantification of DEG expression across endothelial subtypes between WT and miR-125a^Δ/Δ^ cells. **(H)** scRNAseq analysis of endothelial cell subtypes in WT and miR-125a^Δ/Δ^ zebrafish brain. (below) Violin plots of normalized *pgc1a* expression across endothelial subtypes. Two-way ANOVA with Turkey multiple comparisons, p=9.9e-14, 0.9659, 0.0095. **(I)** scRNAseq analysis of human hVOs treated with scramble control or miR-125a inhibitor (miR-125a^inh^), and *PGC1a* expression as in H. Two-way ANOVA with Turkey multiple comparisons, p=6.42e-10, 0.0126, 0.0375, 0.9121. **(J** and **K)** Top ranked Gene Ontology and KEGG pathway enrichment analyses (padj ≤ 0.05) J) in zebrafish and K) in hVOs for miR-125a loss-of-function versus controls **(L)** Mosaic transplantation experiments in zebrafish embryos in which *pgc1a* morpholino (MO)–treated endothelial donor cells (green) were transplanted into WT *kdrl:GFP* hosts (gray). Donor cell localization within PCA/BA and CtA posterior brain vessels was assessed at 32 hpf. **(M)** Quantification of percentage of embryos exhibiting donor endothelial cells at tip cell positions within PCA/BA or CtA territories following *pgc1a* knockdown. Fisher’s exact test, total n= 60 embryos, p=0.0387. **(N)** Endothelial-specific gain-of-function (GOF) via (*fli1a:PGC1a-mcherry*) injection (magenta) into *kdrl:GFP* WT (grey) and assessed in posterior CoW territories relative to CtAs. **(O)** Quantification of the percentage of embryos exhibiting *mcherry+* endothelial cells at tip cell positions within PCA/BA or CtA territories following endothelial-specific *PGC1a* overexpression. Fisher’s exact test, total n=44 embryos, p=0.0012. **(P)** Working model summarizing miR-125a–*pgc1a*–dependent control of endothelial tip-cell specification during posterior brain angiogenesis. High miR-125a levels (blue) constrain *pgc1a* expression (purple) represses CoW tip cell vs CtA specification. All quantifications are presented as mean ± s.e.m. unless otherwise noted. Statistical significance is indicated: ns, not significant, p:: 0.05; *p< 0.05; **p< 0.01; ***p< 0.001; ****p< 0.0001. All data points are reported in Table S1 Sheets 9, 10. Abbreviations as in Fig 3.

*PGC1a* coordinates mitochondrial biogenesis to support cellular metabolic homeostasis in several animal species and cell types, including endothelial cells ^48–50^. Consistent with *pgc1a* de-repression, Gene Ontology and KEGG enrichment analysis of genes upregulated in early miR-125a^Δ/Δ^ brain endothelial cells identified significant overrepresentation of pathways related to glucose oxidation and mitochondrial pathways (**Figures 5C and S6D, Table S1 Sheet 11**).

To assess whether miR-125-derepression of *pgc1a* was specifically linked to brain tip cells, we first performed scRNA-seq on cells isolated from miR-125a^Δ/Δ^ and wild-type brains at ∼26-28 hpf (**Figures S6E-S6H**). Indeed, the most pronounced transcript alterations in miR-125a–deficient cells were confined to the tip cell cluster (**Figures 5D and 5E**), including elevated levels of tip cell–associated genes and an increased proportion of tip cells (**Figures 5F and 5G**). Together, these findings confirm that perturbation of the tip cell population represents the most prominent phenotype resulting from miR-125a loss in zebrafish brain endothelial cells.

Next, we assessed *pgc1a* expression within miR-125a^Δ/Δ^ tip-cell clusters. *pgc1a* mRNA levels were significantly elevated in miR-125a^Δ/Δ^, with tip-cells showing the strongest statistical enrichment (**Figures 5H and S6I**). Importantly, we verified that human *PGC1a* was also de-repressed in miR-125a^inh^ hVO tip cells versus scramble control, with additional, albeit less significant, increases observed in venous endothelial subsets in both species (**Figures 5H and 5I**). Consistently, transcripts enriched in mitochondria function and oxidative phosphorylation also emerged as a top dysregulated pathway in both hVO and zebrafish brain tip cells (**Figures 5J and 5K, Table S1 Sheet 11)**. We infer that de-repression of *PGC1a*, together with activation of metabolic pathways in tip cell populations, is the most conserved signature of miR-125a loss-of-function.

We next asked whether *pgc1a* manipulation is sufficient to specify tip-cell identity in basal CoW and surface CtA brain territories, as predicted if miR-125a acts through *pgc1a* de-repression. To test this, we performed mosaic endothelial cell assays in which *pgc1a* was either suppressed via morpholino injection (before transplantation) or human *PGC1a* was overexpressed under the endothelial *fli1a* promoter ^51,52^. Strikingly*, pgc1a* knockdown conferred tip cell exclusion from CoW territories without affecting CtA integration (**Figures 5L and 5M**). In contrast, mCherry-positive donor endothelial cells overexpressing *PGC1a* were preferentially incorporated into CoW tip cell positions rather than CtA branches (**Figures 5N and 5O**). Notably, parallel *pgc1a* manipulations did not grossly interfere with tip-cell position in the zebrafish trunk region (**Figures S6J and S6K**). Hence, *pgc1a* manipulation phenocopies the brain tip-cell specialization caused by miR-125a loss-of-function and gain-of-function.

These data show that miR-125a tuning of *pgc1a* is a conserved mechanism for brain basal and surface collaterals’ tip cell specification (**Figure 5P**).

### miR-125a Controls Territory-Specific Mitochondrial Function in Sprouting Brain Arteries

Endothelial angiogenic cells are classically viewed as predominantly glycolytic ^53^, yet increasing *in vivo* evidence reveals essential roles for mitochondrial programs in angiogenesis ^54–58^. How metabolic plasticity, particularly mitochondrial regulation, governs the emergence of distinct angiogenic vessels across vascular networks remains unknown. Given our finding that miR-125a and its target *pgc1a* are required for CoW and CtA tip-cell specification, we asked how various mitochondrial features are spatially and temporally regulated during posterior collateral angiogenesis. To address this, we first performed time-lapse imaging of the mitochondrial outer membrane in endothelial cells, using *Tg(fli1a:Tomm20-GFP)* ^59^ in the *Tg(Kdrl:hRASmCherry, fli1a:H2B:BFP)* background to label endothelial cell plasma membranes and nuclei for individual cell analyses (**Figure 6A**).

**Figure 6.**
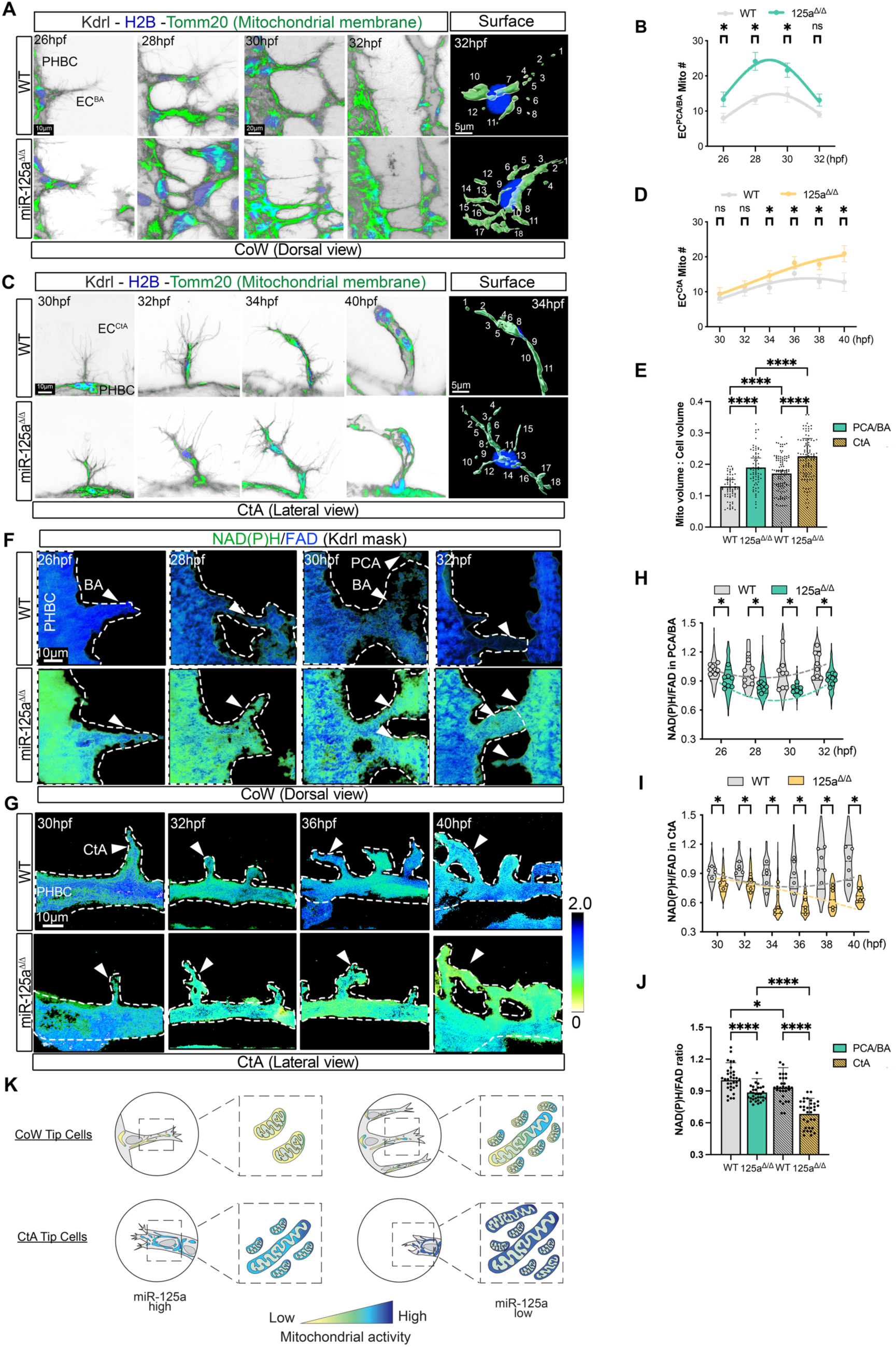
Distinct mitochondrial biogenesis and bioenergetic profiles underlie CoW and CtA angiogenesis and are constrained by miR-125a. **(A)** Representative confocal time-lapse images of mitochondria in endothelial cells during posterior CoW formation (26 to 32 hpf) using *Tg(Kdrl:hRAS-mCH; fli1a:tomm20-eGFP; fli1a:H2B-BFP)* zebrafish in wild-type (WT) and miR-125a^Δ/Δ^ embryos. Nuclei H2B-BFP (blue), mitochondrial membrane Tomm20-eGFP (green), endothelial cell plasma membrane via kdrl (gray). Scale bars as shown. (Right) 3D Imaris surface rendering of segmented mitochondria (green) within individual endothelial cells (blue=nuclei). **(B)** Quantification of mitochondria number per endothelial cell in PCA/BA sprouts from 26 to 32 hpf for WT and miR-125a^Δ/Δ^. Image segmentation and thresholding parameters were applied identically across genotypes using the same analysis pipeline (see STAR Methods). Multiple Mann–Whitney tests, p=0.011, 2.19e-7, 1.57e-3, 0.021, n = 32 endothelial cells per time point per genotype, sampled from 6 embryos each group. See also Fig S6.1. **(C)** Representative confocal live time-lapse images of WT and miR-125a^Δ/Δ^ embryo mitochondria as in A during CtA formation (30-40 hpf). Scale bars as shown. **(D)** Quantification of mitochondrial number per endothelial cell in CtA sprouts from 30-40 hpf as in B. Multiple Mann–Whitney tests, p=0.267, 0.2918, 3.16e-3, 9.46e-3, 2.36e-4, 1.53e-5. n = 20 endothelial cells per time point per genotype, sampled from 6 embryos each group. See also Fig S6.1. **(E)** Ratio of mitochondrial volume to total endothelial cell volume in PCA/BA versus CtA territories for WT and miR-125a^Δ/Δ^. Each dot represents one endothelial cell masked through nuclei H2B-BFP (blue), mitochondrial membrane Tomm20-eGFP (green), endothelial cell plasma membrane vis kdrl (gray) (see STAR Methods). For PCA/BA sprouts, n = 32 endothelial cells per time point per genotype, sampled from 6 embryos each group. For CtA sprouts, n = 20 endothelial cells per time point per genotype, sampled from 6 embryos each group. One-way ANOVA, p=5.84e-77. **(F, G)** Representative two-photon live optical redox imaging of F) CoW vessel angiogenesis (26-32 hpf) and G) CtA vessel angiogenesis (30-40 hpf) in WT and miR-125a^Δ/Δ^ embryos. Arrows=sprouts. White dashed line=vessels. Scale bars shown. **(H)** Quantification of NAD(P)H/FAD ratios from PCA/BA vessels between 26 and 32 hpf across genotypes. NAD(P)H/FAD redox ratios were calculated from endogenous fluorescence using a *kdrl-hRAS-mCherry* mask to delineate artery segments and normalized to WT average redox ratios (see STAR Methods). Each dot represents average redox ratio of an individual vessel (PCA or BA; 2 vessels/embryo) from n= 4 embryos each timepoint, each genotype. Multiple Mann–Whitney tests, p= 0.0053, 0.0034, 0.0117, 0.0135. Violin plot distribution shows regional measurements from within vessels. **(I)** Quantification of NAD(P)H/FAD ratios from CtA vessels between 30 and 40 hpf across genotypes. Each dot represents average redox ratio of one CtA vessel per embryo from n= 5-6 embryos, each timepoint each genotype. Multiple Mann–Whitney tests, p= 0.0159, 0.0043, 0.0303, 0.0159, 0.0043, 0.0022. Violin plot distribution shows regional measurements from within vessels. **(J)** Quantification of average NAD(P)H/FAD redox ratios for PCA/BA versus CtA vessels, across all timepoints in H and I for WT and miR-125a^Δ/Δ^ embryos. Each dot represents average redox ratio per embryo from (PCA/BA) n= 4 embryos each timepoint each genotype, (CtA) n= 5-6 embryos each timepoint each genotype. Ordinary one-way ANOVA with Tukey’s multiple comparisons, p=8.11e-15. **(K)** Working model of mitochondrial biogenesis and activity across both CoW and CtA territories in wild-type and miR-125a^Δ/Δ^. All quantifications are presented as mean ± s.e.m. unless otherwise noted. Statistical significance is indicated: ns, not significant, p:: 0.05; *p< 0.05; **p< 0.01; ***p< 0.001; ****p< 0.0001. All data points are reported in Table S1 Sheets 13, 14. Abbreviations as in Fig 1.

Interestingly, we found that mitochondrial organization differed markedly across brain vascular beds in wild-type embryos: endothelial cells contributing to posterior CoW arteries (PCA/BA) displayed a transient increase in mitochondrial number and volume during active sprouting (26–32 hpf), whereas CtA endothelial cells showed a progressive accumulation of mitochondria from ∼30–40 hpf (**Figures 6A-6D, S7A and S7B**). Consistent with this, CtA endothelial cells exhibited a ∼1.4-fold higher mitochondrial volume compared to PCA/BA cells (**Figure 6E**). Thus, we identify vascular bed–specific mitochondrial phenotypes during brain angiogenesis.

Next, we found that loss of miR-125a led to a global elevation of mitochondrial number and volume in brain endothelial cells across both PCA/BA and CtA territories throughout angiogenic growth (**Figures 6A–E, S7A and S7B**). Despite this overall increase, CtAs remained relatively enriched in mitochondria compared to CoW vessels, indicating that miR-125a modulates mitochondrial abundance without abolishing vascular bed–specific features (**Figure 6E**). Importantly, mitochondrial number and volume ratios were nearly unchanged in trunk intersegmental vessels of miR-125a^Δ/Δ^ embryos (**Figure S7C**), confirming the brain-restricted role for miR-125a.

Together, these findings reveal that mitochondrial content is differentially tuned across distinct cerebrovascular territories and identify miR-125a as a rheostat of such organization.

Elevated mitochondrial biogenesis is associated with elevated cellular energy demand and respiration ^60,61^. To characterize endothelial cell energy and respiration in developing brain vessels, we employed two-photon live optical redox imaging ^62–65^. Optical redox imaging leverages endogenous NAD(P)H and FAD fluorescence, with lower NAD(P)H/FAD ratios as a proxy of increased electron transport–driven oxidation and higher respiration activity (**Figure S7D**). Unexpectedly, endothelial cells sprouting from the PHBC exhibited a lower NAD(P)H/FAD ratio than neighboring neural cells (**Figures S7E and S7F**), indicating a comparatively more oxidized redox state. Strikingly, this metabolic signature was both territory-specific and miR-125a–dependent. In miR-125a^Δ/Δ^ embryos, posterior CoW arteries and CtAs exhibited a significantly reduced NAD(P)H/FAD ratio relative to wild type (**Figures 6F-6I**). This shift was accompanied by a sustained elevation in oxidized FAD levels throughout vessel formation (**Figures S8A–S8C and S9A–S9C)** and aligns with the increased mitochondrial biogenesis and volume observed in mutant endothelial cells.

Notably, these metabolic alterations were confined to brain angiogenic territories: redox ratios were unchanged between wild-type and miR-125a^Δ/Δ^ embryos in trunk ISVs and for the most part in the surrounding brain parenchyma (**Figures S9D-S9F**). Moreover, CtA endothelial cells exhibited a lower NAD(P)H/FAD ratio than CoW endothelial cells in all genotypes, indicating that brain arteries are territorially characterized not only by mitochondrial content, but also by specific mitochondrial bioenergetics (**Figures 6J and 6K**).

Together, these findings reveal that CoW and CtA angiogenesis engage distinct mitochondrial biogenesis and bioenergetic profiles, and that miR-125a functions to constrain metabolic territories.

### miR-125a–*pgc1a*–Dependent ROS Balance Governs Basal and Surface Brain Tip-Cell Behaviors

Despite elevated mitochondrial number and redox state in both miR-125a^Δ/Δ^ CoW and CtA tip cells, CtA segments regress while CoW sprouting persists, resulting in an imbalance in posterior basal and surface endothelial cell distribution and incomplete collateral topologies. How mitochondrial state regulates these distinct angiogenic behaviors is unknown. *PGC1a* couples mitochondrial capacity to redox buffering and maintains reactive oxygen species (ROS) within a growth-permissive range ^66,67^. We hypothesized that *pgc1a*-dependent redox buffering in tip cells explains the distinct sprouting behaviors of CoW and CtAs in miR-125a–mutant embryos. To test this hypothesis, we first analyzed the overall mitochondrial oxidative phosphorylation (OXPHOS) and antioxidant gene expression programs in CoW vs CtA tip cells. *pgc1a*+ tip cells identified in zebrafish brain via scRNA seq were further sub-clustered using CtA-defining markers *mmp25b* and *mmp14* ^28^ to separate CtA versus CoW tip-cell like populations (**Figure 7A**). Distinctly, CtA-tip cells (*pgc1a⁺; mmp25b⁺; mmp14^+^)* had elevated levels of OXPHOS transcripts but reduced induction of antioxidant programs compared with CoW-tip like cells (*pgc1a⁺; mmp25b⁻*; *mmp14 ^−^)* (**Figures 7B, S10A and S10B, Table S1 Sheet 14**). This divergence was markedly amplified in miR-125a^Δ/Δ^ tip cells: OXPHOS pathways increased in both CoW and CtA tip cells, but antioxidant defense was further downregulated in the CtA, but not CoW, tip cells (**Figures 7C, S10A and S10B**).

**Figure 7.**
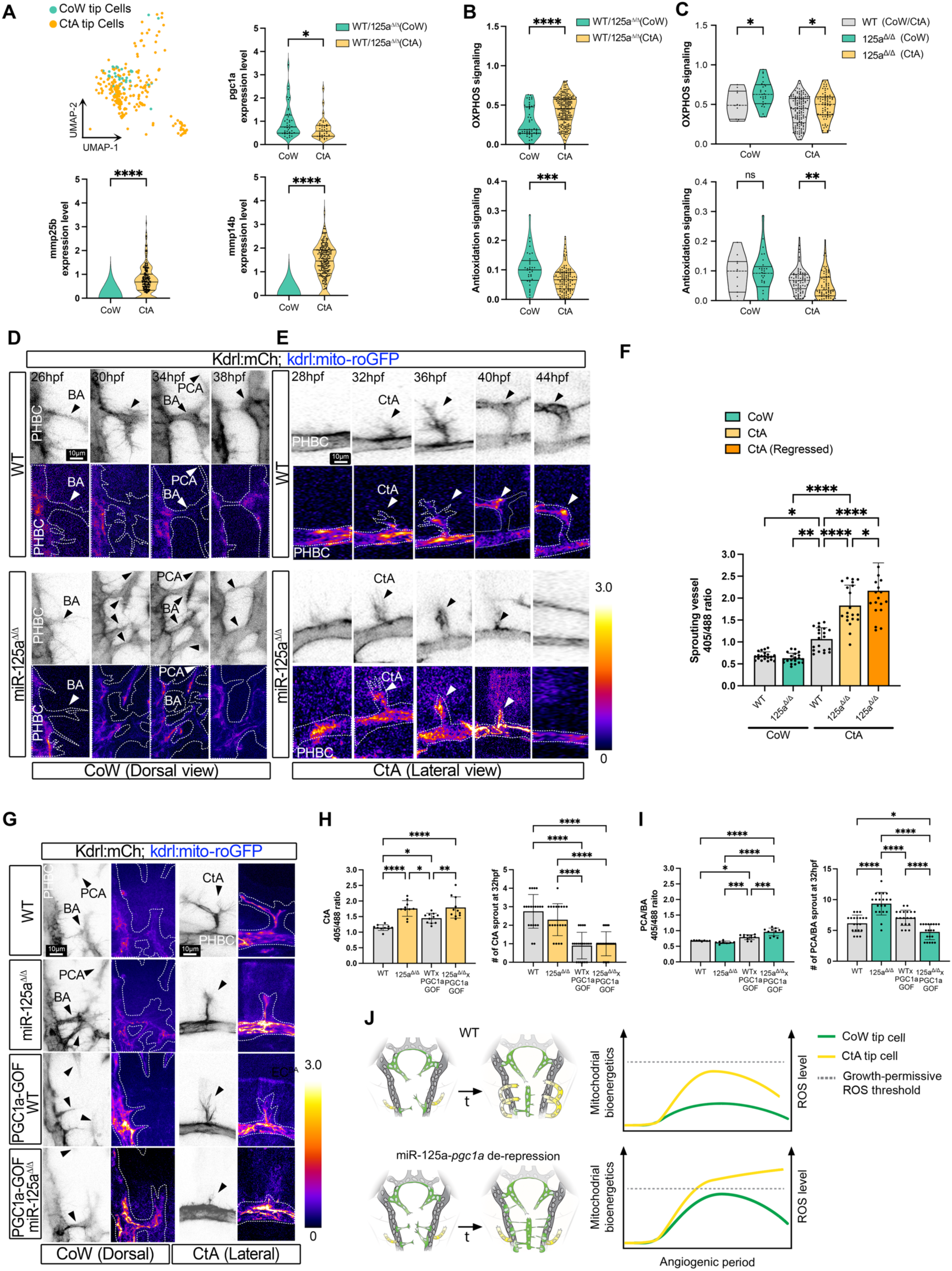
miR-125a–*pgc1a*–Dependent ROS Level Governs Posterior Brain Tip-Cell Behaviors. **(A)** UMAP representation of scRNA-seq data generated from 26-28 hpf miR-125^Δ/Δ^ zebrafish brain endothelial tip cells, subclustered into CoW (green) vs CtA (yellow) regions and violin plots of *pgc1a*, *mmp25b* and *mmp14b* gene expression between the two territories shown. Mann–Whitney test, p=0.0107 (*pgc1a*), p=8.51e-48(*mmp25b*), p=5.84e-77(*mmp14b*). **(B)** Violin plot of *pgc1a*-regulated gene signature scores for OXPHOS and antioxidation pathways in CoW versus CtA tip cells. Gene sets were defined based on *pgc1a*-regulated genes annotated in the STRING network database, and scores were computed using AUCell (AUC-based single-cell gene set scoring). (**C**) Violin plot as in B with gene signature score (AUC-based) in WT vs miR-125a^Δ/Δ^ territories, multiple unpaired t tests. (**D**) Representative confocal time-lapse of *mito-roGFP* in endothelial cells of the PCA/BA during posterior CoW formation (26-38hpf) of WT and miR-125^Δ/Δ^ embryos using the *Tg(Kdrl:hRAS-mCH; kdrl:mito-roGFP)* reporter line (Videos S5 and S6). Dashed line=endothelial cell boundary; Arrow=tip cell. Values shown in Fire lookup table (LUT). Scale bars as shown. (**E**) Representative confocal time-lapse of *mito-roGFP* in endothelial cells of the CtA during formation (28-44hpf) of WT and miR-125^Δ/Δ^ embryos using the *Tg(Kdrl:hRAS-mCH; kdrl:mito-roGFP)* reporter line (Videos S7 and S8). Dashed line=endothelial cell boundary; Arrow=tip cell. Scale bars as shown. **(F)** Quantification of mitochondrial ROS level based on *mito-roGFP sensor* (405/488 ratio) in PCA/BA sprouts (26-38hpf), CtA sprouts and CtA regressed sprouts (28-44hpf) in WT and miR-125a^Δ/Δ^ time-lapses (see STAR Methods). Each dot represents one developing PCA/BA or CtA vessel in WT (n = 4 PCA/BA and 4 CtA time-lapses) and miR-125a^Δ/Δ^ (n = 4 PCA/BA and 6 CtA time-lapses). Ordinary one-way ANOVA test with Tukey’s multiple comparisons, p=4.44e-27. (**G**) Representative confocal live images of mitochondrial ROS level based on *mito-roGFP sensor* (405/488 ratio) in posterior CoW (left) and CtA (right) vessels in WT, miR-125a^Δ/Δ^, *PGC1a*-gain of function (GOF), and 125a^Δ/Δ^;*PGC1a*-GOF embryos at 32hpf. Values shown in Fire lookup table (LUT). Arrows=tip cells. Scale bars as shown. (**H**) Quantification of 405/488 ratio in sprouting CtA vessels (left) and the number of CtA sprouts (right) for all genotypes in G at ∼32hpf. Each dot represents one CtA vessel (1-2 vessels/embryo) from n= 5 WT embryos, 5 miR-125a^Δ/Δ^, 6 *PGC1a*-GOF, and 6 miR-125a^Δ/Δ^;*PGC1a*-GOF. For CtA sprout counting, each dot represents the average for CtA per embryo, n=20 embryos each genotype. One-way ANOVA test, p=3.38e-10(ROS), p=3.94e-12(#). (**I**) Quantification as in H for PCA/BA regions of genotypes in G. For 405/488 ratio quantification, each dot represents one PCA/BA vessel (1-2 vessels/embryo) from n = 4 WT embryos, 4 miR-125a^Δ/Δ^, 6 *PGC1a*-GOF, and 6 miR-125a^Δ/Δ^;*PGC1a*-GOF. For PCA/BA sprouts counting, each dot represents one embryo, n=20 embryos each genotype. One-way ANOVA test, p=2e-15(ROS), p=1.5e-14(#). (**J**) Working model of miR-125a–*pgc1a*–redox regulation within a ROS-growth-permissive range in CoW and CtA endothelial angiogenesis. All quantifications are presented as mean ± s.e.m. unless otherwise noted. Statistical significance is indicated: ns, not significant, p:: 0.05; *p< 0.05; **p< 0.01; ***p< 0.001; ****p< 0.0001. All data points are reported in Table S1 Sheet 15. Abbreviations: CoW, Circle of Willis; PCA, posterior cerebral artery; BA, basilar artery; CtA, central artery; PHBC, primordial hindbrain channel; hpf, hours post-fertilization

These data suggested that elevated mitochondrial activity without a matched antioxidant response could increase mitochondrial H₂O₂ levels and ROS burden specifically in CtA tip cells, leading to angiogenic regression. To test this, we employed the endothelial-specific *Tg(Kdrl:mitochondrial-Rogfp2Orp1)^uto^*^66 68,69^ line, a mitochondrial-targeted ratiometric sensor of H₂O₂–dependent oxidation (**Figures S10C and S10D**). Reporter signals were captured by time-lapse live imaging of posterior CoW and CtA sprouting vessels, co-labeled with *Kdrl:hRAS-mCherry*, between ∼26–44 hpf. In wild-type embryos, mitochondrial ROS signal was significantly higher in CtA tip cells than in CoW tip cells, confirming an intrinsic redox buffer asymmetry between posterior arterial territories at baseline (**Figures 7D-7F, S10E and S10F, Videos S5 and S6**). In miR-125a^Δ/Δ^ embryos, CoW tip cells maintained mitochondrial ROS levels comparable to wild type, whereas CtA tip cells exhibited an ∼2-fold increase relative to controls (**Figures 7D-7F, Videos S7 and S8**). Notably, regressing CtA sprouts displayed higher ROS signals than non-regressing wild-type or mutant counterparts (**Figures 7E and 7F, Video S8**). Together, these data indicate that miR-125a–deficient CtA tip cells undergo disproportionate mitochondrial H₂O₂ accumulation associated with sprout regression.

We next asked whether ROS signaling imbalance and divergent angiogenic behaviors between CoW and CtA territories arise from differences in *pgc1a* activity downstream of miR-125a-regulation. To test this, we quantified ROS signal following endothelia-specific *PGC1a* gain-of-function using a *Tg(fli1a:PGC1a-mCherry)* transgene in wild-type or in miR-125a^Δ/Δ^, in which *pgc1a* is de-repressed at baseline (**Figures 7G-7I and S6K**). Endothelial *PGC1a* overexpression in wild-type was sufficient to recapitulate the CoW and CtA mitochondria ROS profile and angiogenic phenotypes observed in miR-125a–deficient embryos (**Figures 7G-7I and S10G**). *fli1a:PGC1a* embryos developed into larval and adult stages and exhibited an increased frequency of incomplete CoW and decreased SBC richness, consistent with those observed in adult miR-125a^Δ/Δ^ animals (**Figures S10H and S10I**). Next, we assessed the effect of endothelia *PGC1a* overexpression in miR-125a mutants. Notably, miR-125a^Δ/Δ^; *fli1a:PGC1a* CtAs did not further increase the already elevated ROS signaling in CtA tip cells; however, the frequency of regressing CtA segments was increased (**Figures 7G, 7H and S10G**). In contrast, in the same embryos, CoW tip cells displayed a quantifiable increase in mitochondrial ROS levels, and consistent with this elevated oxidative burden, we observed fewer CoW angiogenic sprouts in miR-125a^Δ/Δ^; *fli1a:PGC1a* embryos compared with miR-125a^Δ/Δ^ (**Figures 7G, 7I and S10G**).

Thus, miR-125a–*PGC1a* signaling sets mitochondrial ROS levels that instruct basal and surface brain tip-cell behavior to encode brain collateral architecture (**Figure 7J**).

### Restoration of Angiogenic Behavior and Posterior Collateral Architecture by *pgc1a*-Dependent Metabolic Rebalancing in the miR-125a loss-of-function model

Our data suggest that loss of miR-125a–*PGC1a* dependent repression alters mitochondrial metabolism in embryonic tip cells, leading to incomplete posterior primary and secondary collateral network topologies and less brain vascular resilience. To test whether *pgc1a* could be an entry point to correct collateral growth back to completeness in miR-125a^Δ/Δ^ mutant models, we crossed miR-125a^Δ/Δ^ mutants with adult-viable *pgc1a* homozygous mutants (*ppargc1a^bns176 70,71^*) to obtain miR-125a^+/Δ^; *pgc1a^+/−^* double heterozygotes. We quantified mitochondrial bioenergetic states, sprouting behaviors, and collateral architecture at larval and adult stages, comparing them with miR-125a^+/Δ^ and wild-type controls (**Figure 8A**).

**Figure 8.**
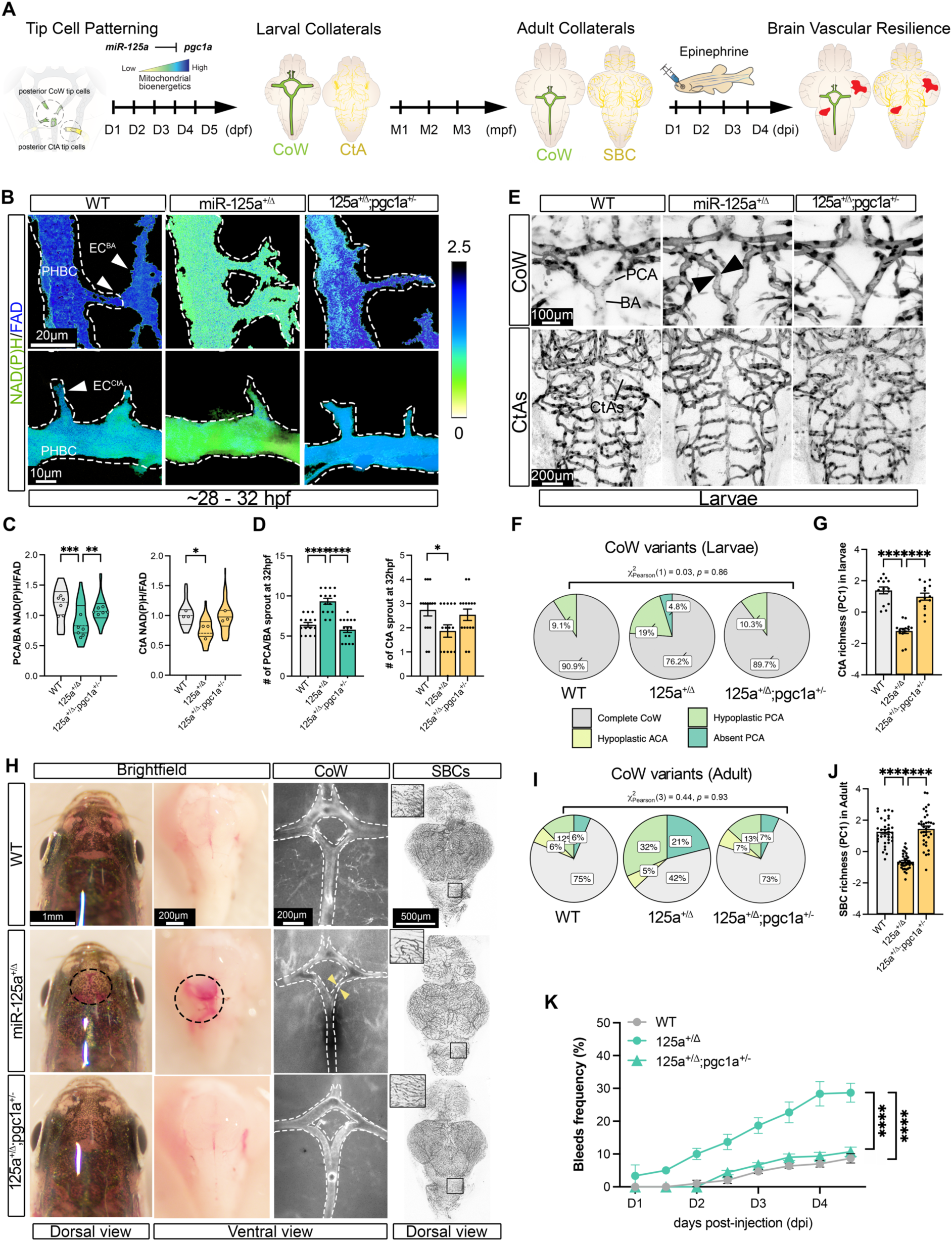
miR-125a–*pgc1a*–dependent mitochondrial regulation links developing collateral topologies to adult brain vascular resilience. **(A)** Experimental schematic of miR-125a-*pgc1a* genetic complementation assay to rebalance mitochondrial metabolism and posterior brain angiogenic output and adult brain vascular resilience. Zebrafish miR-125a^Δ/Δ^ were crossed with *pgc1a^-/-^ (ppargc1a^bns176^*) to generate double heterozygotes (miR-125a^+/Δ^; *pgc1a^+/−^*). Tip cell bioenergetics and sprouting were assessed at 32 hpf, CoW and CtA anatomies were assessed in larvae after 5 days (D), and CoW and SBC collateral variations were assessed in adults after ∼3 months (M) with functional cerebrovascular resilience tested by 4 days of epinephrine-driven hemodynamic stress. **(B)** Representative two-photon optical redox images of posterior brain vessels showing NAD(P)H/FAD ratios in CoW PCA/BA and CtA vessels of WT, miR-125a^+/Δ^, miR-125a^+/Δ^;*pgc1a^+/−^*embryos. Dashed line=vessels. Scale bars as shown. **(C)** Quantification of NAD(P)H/FAD ratio from PCA/BA (left) and CtA (right) vessels across genotypes shown in B. Each dot represents average redox ratio of PCA and BA (2 vessels/per embryo) or CtA vessel from n= 3 embryos each genotype. One-way ANOVA test, p= 9.733e-5, 0.0272. Violin plot distribution shows regional measurements from within vessels. **(D)** Quantification of sprout number in PCA/BA vessels (left) and CtA (right) across genotypes shown in B. Each dot represents one embryo from (PCA/BA) n=15 embryos each genotype and (CtA) n=15 each genotype. Ordinary one-way ANOVA, p= 6.044e-9, 0.0436. **(E)** Representative confocal images of larval posterior CoW variants (top) and CtA networks (bottom) visualized in *Tg(kdrl:GFP)* fish. Arrows=hypoplastic vessel. Scale bars as shown. **(F)** Distribution of larval CoW anatomical variants. No significant difference between WT and double heterozygotes (miR-125a^+/Δ^; *pgc1a^+/−^)* in larval CoW configuration (χ²(1) = 0.03, p = 0.86 ns, Cramer’s V=0.00) additional comparisons in S8.1C. **(G)** Quantification of CtA richness in larvae. Each dot represents one larva, n=14 larvae each genotype. Ordinary one-way ANOVA test, p=3.75e-11. **(H)** Representative dorsal brightfield images of adult heads (left), alongside brightfield dissected brain base (middle) and the corresponding fluorescent images of CoW configuration (right, *kdrl:GFP*) in ventral view and SBCs network in dorsal view. Dashed circle=bleed event. Dashed line = CoW vessels. Arrow=hypoplastic arteries. Scale bars as indicated. **(I)** Distribution of adult CoW anatomical variants. No significant difference between WT and double heterozygotes (miR-125a^+/Δ^; *pgc1a^+/−^)* in adult CoW configuration (χ²(3) = 0.44, p = 0.93 ns, Cramer’s V=0.00), with Cramer’s V reported to measure strength of association independent of sample size. Additional comparisons in S8.1E. **(J)** Quantification of adult SBC richness. Each dot represents one adult from n= 21 fish each group. Ordinary one-way ANOVA test, p=0.0017. **(K)** Quantification of brain bleed frequency. Each group consisted of 30 fish from three independent experiments, and dots are average over all experiments. Ordinary two-way ANOVA test at D4 (p=2.79e-11 (WTvs125a^+/Δ^), 0.68 (WTvs125a^+/Δ^;*pgc1a^+/-^*), 8.75e-10 (125a^+/Δ^vs125a^+/Δ^;*pgc1a^+/-^*). All quantifications are presented as mean ± s.e.m. unless otherwise noted. Statistical significance is indicated: ns, not significant, p:: 0.05 (not shown); *p< 0.05; **p< 0.01; ***p< 0.001; ****p< 0.0001. All data points are reported in Table S1 Sheets 16, 17. Abbreviations: CoW, Circle of Willis; SBCs, surface brain collaterals; PCA, posterior cerebral artery; BA, basilar artery; CtA, central artery; PHBC, primordial hindbrain channel; hpf, hours post-fertilization

First, optical NAD(P)H/FAD redox imaging during CoW and CtA sprouting revealed that miR-125a^+/Δ^ embryos had a lower NAD(P)H/FAD ratio than wild-type embryos, indicating that partial loss of miR-125a is sufficient to induce FAD oxidation (**Figures 8B, 8C, S11A and S11B**). Importantly, NAD(P)H/FAD ratios in miR-125a^+/Δ^;*pgc1a^+/−^* endothelial cells were restored to wild-type levels in both CtA and CoW territories (**Figures 8B, 8C, S11A and S11B**), coinciding with normalization of CtA and BA/PCA sprouting segments (**Figure 8D**). Thus, partial reduction of *pgc1a* is sufficient to restore CoW and CtA territorial mitochondrial bioenergetics and angiogenic output in the setting of miR-125a haploinsufficiency.

Consistent with these findings, miR-125a^+/Δ^; *pgc1a^+/−^*larvae displayed normalized CoW configuration and CtA richness (**Figures 8E–8G, S11C and S11D**). Notably, adult miR-125a^+/Δ^; *pgc1a^+/−^*animals showed robust rescue of complete posterior collateral anatomy compared to miR-125a^+/Δ^ alone (**Figures 8H–8J, S11E and S11F**).

Given that low miR-125a levels are associated with incomplete posterior CoW anatomy and increased vulnerability to vascular injury in humans, paralleling the miR-125a^+/Δ^ zebrafish phenotype (**Figures 1N–1P**), we next asked whether *pgc1a* complementation could also restore functional cerebrovascular resilience. Indeed, loss of a single *pgc1a* allele in the miR-125a^Δ/+^ background reduced bleeding incidence in miR-125a^+/Δ^ to wild-type levels (**Figures 8H and 8K**). Thus, we demonstrate that reduced *pgc1a* dosage resolves brain vascular vulnerability caused by miR-125a deficiency.

Together, these findings show that tuning *pgc1a* levels in a miR-125a partial loss model restores a completely connected primary basal and richly branched secondary surface collateral architectures and confers lifelong cerebrovascular resilience.

## Discussion

Here, we identify mitochondrial metabolism as a “tip-cell patterning” mechanism that establishes multiscale brain collateral architecture. Through integrated genetic, developmental, and functional analyses in zebrafish, supported by cross-species evidence, we define a miR-125a–*pgc1a* axis that tunes mitochondrial biogenesis and redox buffering to establish territory-specific mitochondrial states in brain tip cells. We propose that sprouting angiogenesis requires a finely titrated increases in mitochondrial output, and that miR-125a-mediated restraint of *pgc1a* activity maintains this output within a growth-permissive redox range. Loss of miR-125a disrupts this balance, producing region-specific oxidative burden that differentially rewires tip-cell behaviors in CoW versus CtA territories. Strikingly, these metabolic perturbations do not simply alter angiogenic growth; rather, they direct new vascular patterning, generating collateral networks with distinct incomplete geometry and poor connectivity that persist into adulthood and compromise resilience under hemodynamic stress. Together, our findings reveal that the basal and surface vascular network architecture is not a byproduct of broad angiogenesis, but is encoded by a tip-cell program driven by mitochondrial metabolism, providing a new entry point for understanding and potentially restoring brain vascular resilience.

The collateralization of the brain has long been recognized for its remarkable anatomical hierarchy. Previous studies have shown that genetic or environmental factors can independently influence the formation of primary and secondary collateral networks in the brain ^24–26,72–75^. As a result, most of our knowledge about collateral formation comes from studies focused on a single vascular bed. Thus, it is unclear whether brain collateral networks are governed by shared biological mechanisms or whether network-specific programs account for the substantial variation seen among human population ^14,23,72,73,76–78^. Our data show that the basal CoW and surface pial collateral networks arise from a pre-organized developmental template under the guidance of molecular rheostats, such as microRNAs that titrate patterning programs. MicroRNAs represent particularly effective rheostats for generating inter-individual variability, because their regulatory effects can be sensitive to quantitative fluctuations in target transcript abundance, timing, and translation dynamics ^79,80^, parameters that may amplify biological variability during a narrow developmental window. Our data also support that anterior brain collateralization may be governed by distinct pathways, as miR-125a-dependent regulation is required for the formation of complete and rich posterior, but not anterior, collateral patterning. A limitation of our zebrafish analyses is that subtle anterior variants (e.g., AComA configuration) are difficult to quantify reliably in dissected brains, motivating future work using improved imaging strategies in intact animals. Nonetheless, no anterior SBC variation was associated with CoW variants (anterior or posterior) in our zebrafish data, and human radiographic cohort analyses converge on a posterior bias, supporting a miR-125a-specific posterior patterning program.

It is well established that anterior and posterior brain vessels differ in their developmental timing and regulatory programs ^42,81^, in agreement with the either/or presentation of human anterior and posterior collateral variants ^82^. Hence, the specialization of miR-125a-*pgc1a* regulation in posterior and not anterior variants is consistent with differences in developmental signaling and timing that characterize the development of these vessels. Differences in vascular maturation at later developmental stages, including earlier acquisition of smooth muscle coverage in anterior segments, may further contribute to this regional selectivity ^83,84^. Finally, our analyses of cerebrovascular resilience under hemodynamic stress did not detect region-specific bleeding events in miR-125a loss-of-function models (e.g., anterior vs posterior brain regions); instead, vascular injury was distributed across the brain. We speculate that this may reflect system-level hemodynamic consequences, such as altered red blood cell flux and flow instability within a structurally disorganized network, rather than failure of a single vulnerable territory. Furthermore, under stress conditions, collateral pathways throughout the brain are remodeled to preserve perfusion, which may further blur the anatomical localization of vascular injury. Overall, miR-125a loss affects global brain vascular resilience, but this vulnerability can be prevented by correcting downstream signaling, positioning miR-125a as a tractable entry point for improving brain health.

In the human population, anatomical variation in the CoW has been associated with increased risk of damaging cerebrovascular events ^14,23,77,85,86^. Here, leveraging a cohort enriched for individuals with radiographic evidence of covert brain injury ^38^, we identify significantly lower circulating miR-125a levels in participants with posterior, but not anterior incomplete CoW anatomies. In humans, pial collateral vessel density cannot be measured as directly or as precisely as in animal models. For example, the MRA methodology used here to visualize primary collaterals does not reliably resolve small-diameter vessels. Consequently, this approach cannot accurately capture both primary and secondary collateral vessel density. Another important caveat is that our cohort size is limited, which reduces statistical power to detect miR-125a–associated genetic effects (miR-eQTLs) and prevents us from performing robust causal inference analyses. Consistent with this limitation, our Mendelian randomization analysis showed directionally positive trends linking genetic predictors of miR-125a levels to posterior (but not anterior) CoW variation, but these associations did not reach statistical significance (data not shown). Larger cohorts in combination with future sophisticated cerebrovascular human imaging analyses will be required to test whether miR-125a regulation causally contributes to both posterior primary and secondary collateral configurations.

Nevertheless, circulating miR-125a levels positively correlated with complete collateral architectures. Notably, miR-125a has recently been identified as a blood-based biomarker linked to cognitive domains, brain regions, and neuronal processes affected by aging and neurodegeneration, traits that overlap with features of covert brain injury ^87,88^. More broadly, circulating miRNAs are emerging as robust biomarkers across diverse human diseases ^35–37^, in part because they are unusually stable in plasma/serum, being protected from RNase degradation through association with extracellular vesicles and non-vesicular carriers (e.g., Argonaute2 protein complexes and lipoproteins ^89,90^). Circulating miRNAs can reflect tissue injury and cellular turnover, but may also be actively released through regulated secretion, potentially supporting intercellular communication. Although the mechanisms underlying reduced circulating miR-125a remain unresolved, our findings position miR-125a as an attractive candidate biomarker for CoW collateral anatomies associated with vascular vulnerability that may translate into long-term risk.

We show that brain endothelial tip cells are surprisingly weakly glycolytic, yet exhibit elevated mitochondrial biosynthesis and high bioenergetic capacity throughout angiogenesis. Moreover, tip cells from distinct brain vascular territories are differentially wired to mitochondrial programs that tune oxidative phosphorylation versus redox buffering, and these metabolic configurations have functional consequences for angiogenic behavior. Together, our findings reveal that tip-cell function depends on this precisely regulated mitochondrial state. This model contrasts with the prevailing view that tip cells rely predominantly on glycolysis to support rapid migration and sprouting ^53,91–93^. However, tip cells are highly heterogeneous and not functionally interchangeable, making it unlikely that a single energetic strategy uniformly supports all tip-cell states across tissues and vascular territories. Recent work using human *in vitro* sprouting assays has shown that tip cells can adopt flexible metabolic programs, including reduced reliance on glycolysis relative to neighboring stalk cells ^94^. While our data do not exclude an essential contribution of glycolytic pathways to angiogenesis, or account for potential differences in total co-enzyme pool size, we uncover a tip-cell, territory-specific preference for mitochondrial metabolism that shapes tissue-specific angiogenic behaviors. In our model, this metabolic state is controlled by miR-125a through post-transcriptional regulation of *pgc1a*. Because *pgc1a* can rapidly coordinate multiple metabolic outputs—including mitochondrial biogenesis, respiratory capacity, and redox balance—it represents an ideal regulatory node to couple mitochondria metabolic plasticity and tip-cell specialization within the narrow temporal window of angiogenic decision-making.

Molecularly, *PGC1a* is an inducible transcriptional co-factor that orchestrates broad metabolic remodeling, including mitochondrial biogenesis, oxidative phosphorylation, and glucose/fatty acid metabolism—programs that can support tissue growth and adaptation during development and stress ^51,95^. In cardiomyocytes, *Pgc1a* overexpression has shown to promote mitochondrial biogenesis and lead to cardiomyopathy ^96^. Importantly, *PGC1a* has also been directly implicated in angiogenesis, although its functional directionality appears highly context-dependent. In some settings, *PGC1a* induces *VEGFA* expression (often through ERRα-dependent transcriptional programs) and promotes angiogenesis independently of canonical HIF signaling, consistent with a role in coordinating oxygen delivery with oxidative metabolic demands ^97^. Conversely, endothelial *PGC1a* has been reported to inhibit endothelial migration and sprouting by activating Notch signaling, contributing to vascular dysfunction in metabolic disease contexts ^98^. Together, these findings suggest that *PGC1a* does not act as a uniform pro-or anti-angiogenic factor, but rather functions as a tunable metabolic node whose impact depends on cellular state, tissue context, and redox constraints. Consistent with this view, *PGC1a* can bi-directionally shape ROS biology by simultaneously increasing respiratory capacity (and therefore oxidative load) while also inducing antioxidant defense programs that buffer ROS and maintain cellular function ^49^. We propose that this dual energy vs ROS-tuning capability underlies the divergent angiogenic outcomes across brain artery territories. In miR-125a–deficient embryos, CtA tip cells exhibit increased oxidative burden together with weaker antioxidant buffering relative to CoW tip cells, culminating in sprout regression. Notably, forced *PGC1a* expression in CoW tip cells also induces regression, supporting a model in which excessive mitochondrial output—when uncoupled from adequate buffering—destabilizes persistent sprouting. Thus, we propose that miR-125a with other territory-specific niche cues (e.g., cell identity, oxygen gradients, and upstream signaling inputs) calibrate *PGC1a* -dependent mitochondrial output and antioxidant capacity to enable tissue-specific tip cell behaviors.

Overall, our findings establish how the entire brain vascular network architecture is patterned through metabolic specialization of endothelial tip cells. By identifying miR-125a–*pgc1a* control of mitochondrial output and redox buffering as determinants of territory-specific tip-cell behavior, we link the cell-intrinsic bioenergetic state to brain network geometry, long-term stability, and collateral resilience. Since collaterals are a lifesaving vascular structure, they are organized hierarchically and interconnect across multiple vascular scales in many organs. Thus, our work raises the exciting possibility that organ-specific collateral architectures are broadly encoded by pool of embryonic cells via conserved metabolic patterning programs, and may even be reflected by collateral-specific circulating biomarkers. Defining these programs across organs could reveal general principles of collateral network design and provide new routes to quantify vulnerability and resilience across organ systems.

